# CryoEM Structures of the Human HIV-1 Restriction Factor SERINC3 and Function as a Lipid Transporter

**DOI:** 10.1101/2022.07.06.498924

**Authors:** Susan A. Leonhardt, Michael D. Purdy, Jonathan R. Grover, Ziwei Yang, Sandra Poulos, William E. McIntire, Elizabeth A. Tatham, Satchal Erramilli, Kamil Nosol, Kin Kui Lai, Shilei Ding, Maolin Lu, Pradeep D. Uchil, Andrés Finzi, Alan Rein, Anthony A. Kossiakoff, Walther Mothes, Mark Yeager

## Abstract

The host proteins SERINC3 and SERINC5 are HIV-1 restriction factors that reduce infectivity when incorporated into the viral envelope. The HIV-1 accessory protein Nef abrogates incorporation of SERINCs via binding to intracellular loop 4 (ICL4). CryoEM maps of full-length human SERINC3 and an ICL4 deletion construct reveal that hSERINC3 is comprised of two *α*- helical bundles connected by a ∼40-residue, tilted, “crossmember” helix. The design resembles non-ATP-dependent lipid transporters. Consistently, purified hSERINCs reconstituted into proteoliposomes flip phosphatidylserine (PS), phosphatidylethanolamine and phosphatidylcholine. SERINC3 and SERINC5 reduce infectivity and expose PS on the surface of HIV-1 and also MLV, which is counteracted by Nef and GlycoGag, respectively. Antiviral activities by SERINCs and the scramblase TMEM16F correlate with the exposure of PS and with altered conformation of the envelope glycoprotein. We conclude that SERINCs are lipid transporters, and we demonstrate that lipid flipping is directly correlated with loss of infectivity.

**One Sentence Summary:** The HIV-1 restriction factor SERINC3 has a molecular design similar to non-ATP dependent lipid transporters, a function supported by the observation of flipping activity in proteoliposomes and exposure of phosphatidylserine on HIV-1 and MLV particles, which is correlated with loss of infectivity.

---

SERINC proteins are comprised of a family of five isoforms with 31-58% amino-acid sequence identity, which are thought to incorporate serine into phospholipids^1^. The ∼50 kDa integral membrane proteins have ten predicted transmembrane domains and a single N-glycosylation site **(Extended Data Fig. 1a,c)**. In 2015, substantial interest in SERINCs was stimulated by the observation that the presence of host proteins SERINC3 or SERINC5 in the envelopes of HIV-1 particles reduced infectivity^2, 3^. The restriction activity was highest for SERINC5, followed by SERINC3, and was not detected for SERINC2. The restriction activity of SERINC5 was counteracted by the viral protein Nef, which redirected SERINC5 to an endosomal compartment, thereby precluding incorporation into the viral envelope^4^. In addition to HIV-1, restriction activity of SERINC5 has been observed for other enveloped viruses including murine leukemia viruses (MLV) that express GlycoGag to counteract SERINCs^2, 3, 5, 6^.

SERINCs may have multiple functions that collectively interfere with virus entry. SERINC incorporation into HIV-1 particles appears to affect the conformation of the envelope glycoprotein (Env) as measured by increased exposure of sequestered epitopes, and it interferes with membrane fusion by impeding Env clustering, intermediates in the fusion pathway^7^ and expansion of the fusion pore^8–11^. Although SERINCs may impede fusion by effects on the local lipid composition in the envelope, lipid quantitation and fractionation analysis have shown no changes in the amount and content of cell and viral membrane lipids in the presence of SERINC5^12^. Given this background of mechanistic uncertainty, we sought to determine high-resolution structures of a human SERINC protein and ascertain the the mechanism of its antiviral activity.

## Structure determination and molecular design

We tested a variety of recombinant systems, and the protein yield was highest for human SERINC3 expressed in Sf9 insect cells **(Extended Data Fig. 2 a,b)**. Formation of a complex with a synthetic Fab **(Extended Data Fig. 2c-i)** enabled determination of a map of the 50 kDa, full-length, wild type (WT) hSERINC3 monomer by the use of cryoEM and single-particle image analysis (**Fig. 1a,b, Extended Data Figs. 3, 4**). Göttlinger and colleagues presented evidence that the ability of Nef to downregulate hSERINC5 involves ICL4^4^, which contains an acidic cluster motif (EDTEE) that is a proposed binding site for the clathrin adapter protein, AP2^13^ (Comparable residues in SERINC3 are SDEED, **Extended Data Fig. 1b**). Hurley and colleagues demonstrated that AP2 can form a stable complex with Nef^14^, and a hSERINC5-AP2-Nef complex may provide a mechanism for targeting of hSERINC5 to the endosomal/lysosomal pathway in Nef-containing HIV-1 strains^2, 3, 15^, thereby preventing restriction by abrogating incorporation of hSERINC5 into the HIV-1 envelope. For these reasons, we also generated a cryoEM map of the ICL4 deletion mutant (*Δ*ICL4) of hSERINC3 (**Fig. 1c,d, Extended Data Figs. 3 and 5**).

**Figure 1.**
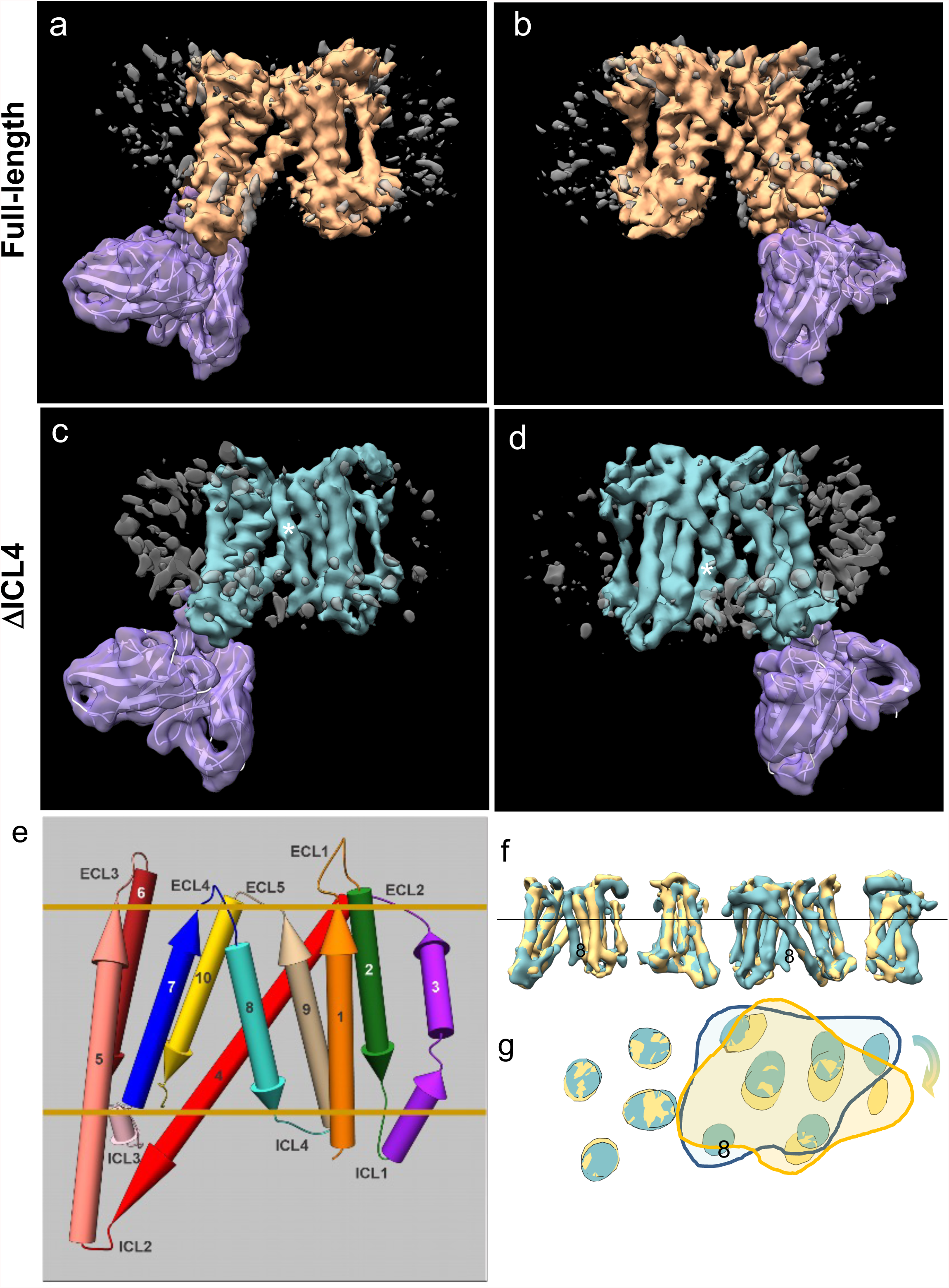
CryoEM density maps of full-length, WT hSERINC3. **(a, b; gold)** and a deletion mutant of the fourth intracellular loop (*Δ*ICL4) **(c, d; blue)** and the variable region of the bound Fab (purple). **(e)** The secondary structure model shows that hSERINC3 is comprised of two *α*-helical bundles. The proximal Fab-binding bundle contains H5, 6, 7 and 10, and the distal bundle contains H1, 2, 3 and 9. The two bundles are connected by a long 40 residue, diagonal “crossmember” *α*-helix (H4). H4 is paired with H8, which displays conformational variability in the full-length WT map and is better-ordered in the *Δ*ICL4 deletion mutant (asterisks in (c) and (d). **(f,g)** Superposition of the cryoEM density maps of full-length, WT hSERINC3 (gold) and the *Δ*ICL4 deletion mutant (blue). (f) Horizontal line in the cross-section view identifies the position of the superimposed *en face* view in (g). The *Δ*ICL4 deletion elicits rotation of the less ordered bundle.

The maps of WT and *Δ*ICL4 hSERINC3 are comprised of two *α*-helical bundles connected by a ∼40-residue, highly-tilted, “crossmember” *α*-helix (H4) **(Figs. 1, 2)**. The overall resolution of the WT hSERINC3 map was 4.2 Å (**Extended Data Fig. 4d**). However, the resolution of the Fab-binding bundle (Fab-proximal, H5,6,7,10) was higher (∼3.6 Å), with clear definition of several tyrosine and tryptophan residues in the Fab CDR loops and the hSERINC3 epitope **(Fig. 2)**. The map in the region of the distal *α*-helical bundle (H1,2,3,9) has a lower resolution (∼4.4 Å) due to conformational variability between the domains (**Fig. 2a,b**). The hSERINC3 maps recapitulate the high-resolution cryoEM structure of a Drosophila SERINC ortholog (TMS1d) (**Extended data Fig. 6a,b,d,e**)^16^. Given that the sequence identities between hSERINC3 and hSERINC5 and hSERINC2 are 39.1 and 51.9% (**Extended Data Fig. 1c**), we expect that the molecular design will be conserved in the SERINC family (**Fig. 1e**). This inference is supported by the 7.1 Å cryoEM map of hSERINC5, which shows rod-like densities consistent with *α*-helices in a similar disposition with the *α*-helical bundles visualized in hSERINC3^16^. Furthermore, AlphaFold predicts conservation in the transmembrane architecture of SERINC3, SERINC5 and SERINC2 (**Extended Data Fig. 6g-j**).

**Figure 2.**
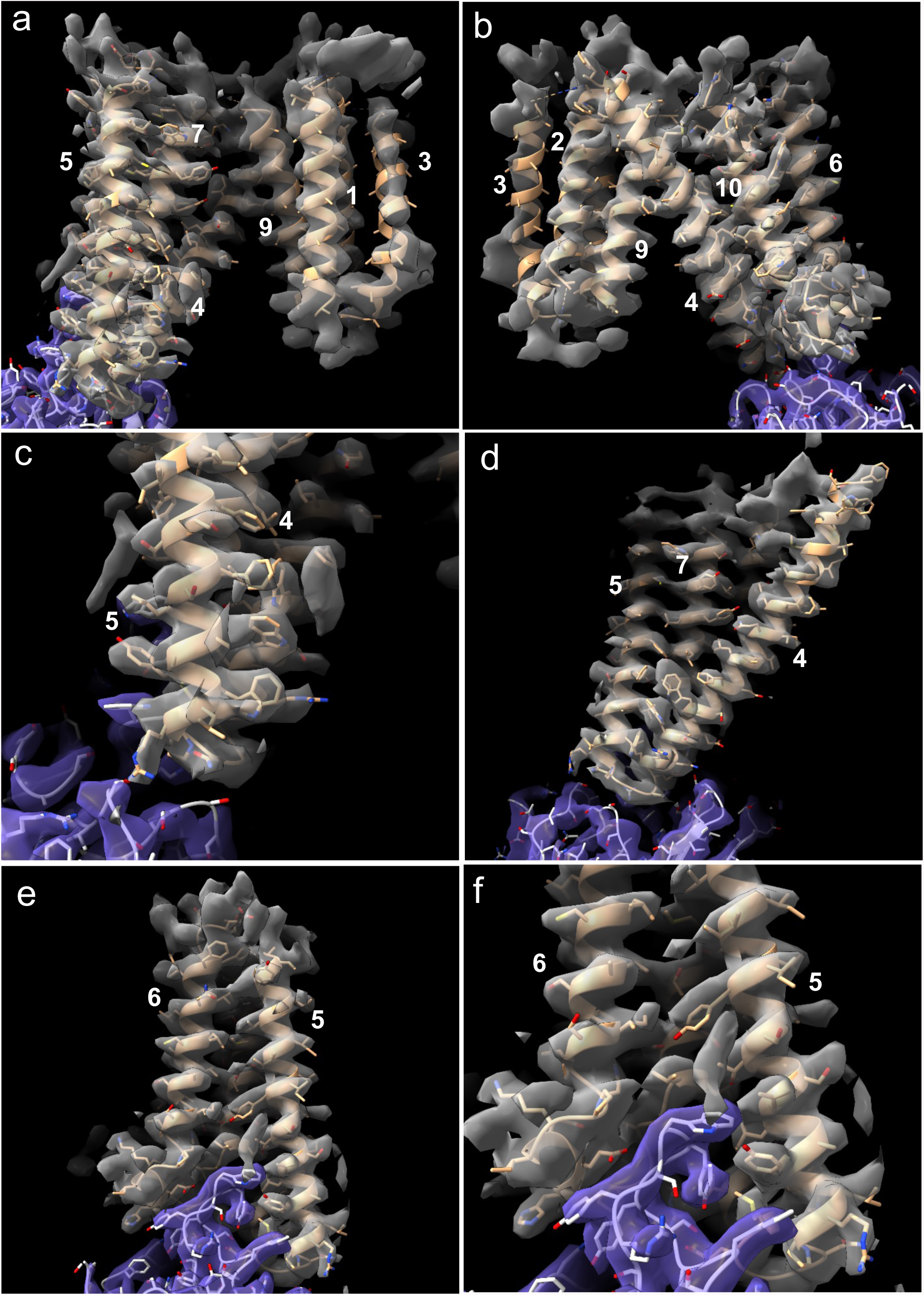
CryoEM density map and refined structure of WT hSERINC3-Fab complexes (*α*- helices labeled as in. Figure 1e). CryoEM density map of WT hSERINC3 (gold) and the variable region of the bound Fab (purple). The cryoEM density map is shown as transparent white (hSERINC3) or transparent purple (Fab), with the fitted model of hSERINC3 (cartoon colored in gold) and Fab (white cartoon). (**a,c**) Map displays two *α*-helical bundles connected by a highly tilted 40 residue “crossmember” helix. (**a,b**, related by a 180° rotation) (**b**) Counterclockwise rotation by 90° about the vertical and enlargement of the Fab-binding helical bundle from the position in (**a**), emphasizing an oblong density with a cluster of adjacent tyrosine residues. (**d**) Counterclockwise rotation by 90° about the vertical from the position in (**b**) showing the highly tilted 40 residue “crossmember” helix. (**e**) View highlighting the Fab variable region where the CDR loops bind to an hSERINC3 conformational epitope. (**f**) Enlargement of (**e**) showing the aromatic side chains in the CDR loops. The map resolution of the Fab-proximal bundle was ∼3.6 Å, with well-defined side chain densities. The distal bundle has a resolution of ∼4.4 Å, with well- defined bulky side chains.

Using the Fab-proximal bundle as a reference, the distal bundle is rotated ∼5° in the *Δ*ICL4 map (**Fig. 1f, g**) relative to the conformation in WT hSERINC3. Interestingly, the density for the H8 helix was ill-defined in the WT hSERINC3 cryoEM map **(Fig. 1a,b)**, whereas H8 was well-defined in the *Δ*ICL4 map and in TMS1d ^16^ (**Fig. 1c,d**, asterisk). We infer that the ordering of H8 and the conformational change between the helical bundles in the *Δ*ICL4 mutant suggest that there may be allosteric communication between ICL4 and the transmembrane *α*-helices.

## Structural similarity of SERINC proteins with non-ATP-dependent, unregulated lipid transporters

ATP-dependent lipid transporters contain ATP-binding cassettes and are categorized as outer to inward flipping P4-type ATPases or “flippases” and inward to outward flipping ABC transporters or “floppases”^17–20^. SERINCs clearly do not have ATP-binding cassettes (**Extended Data Fig. 1a**). Non-ATP-dependent or unregulated lipid flipping proteins are designated as “scramblases” and exhibit both inward to outward and outward to inward lipid flipping. We were intrigued that hSERINC3 bears a structural resemblance with unregulated lipid transporters such as archaeal PfMATE^21^ and bacterial proteins MurJ^22, 23^(**Extended Data Fig. 1d-f**) and LtaA^24^, which are comprised of two *α*-helical bundles connected by a long, highly-tilted crossmember *α*-helix. As has been proposed for the clefts between the helical bundles of PfMATE^21^, MurJ^22, 23^ and LtaA^24^, we speculate that the cleft between the helical bundles in hSERINC3 may serve as an entry portal for lipids. Consequently, we infer that all three SERINC isoforms may function as non-ATP-dependent, unregulated lipid transporters.

## Purified hSERINC3 reconstituted into proteoliposomes induces flipping of phosphatidylcholine, phosphatidylethanolamine and phosphatidylserine

The most compelling evidence for lipid transport activity is provided by analysis of liposomes assembled from chemically defined synthetic lipids, with or without reconstitution of the putative lipid transporter. Notably, phospholipids display minimal spontaneous flipping activity in liposomes lacking lipid transporters, water pores, detergents or surfactants^19, 25, 26^. The experimental design to measure lipid flipping is depicted in **Fig. 3a** and is based on the work of Menon and colleagues^27^. A critically important control is to demonstrate that incorporation of the lipid transporter does not result in nonspecific leakiness of the liposomes (**Fig. 3a, ii**). This is accomplished by trapping water soluble NBD-glucose in the liposomes containing the putative lipid transporter. In generating the liposomes, some residual NBD-glucose resides in the extravesicular space, which accounts for an initial reduction in the fluorescent signal upon addition of dithionite, a membrane impermeable reducing agent (**Fig. 3b**, 100 sec time point). Importantly, the fluorescent signal reaches a stable plateau for liposomes containing a high concentration (1.5 µg/mg lipid) of hSERINC3 (**Fig. 3b**, 100-500 sec). If the proteoliposomes had been leaky, then the signal would have continued to decline during extended incubation. Lastly, addition of Triton X100 solubilizes the liposomes, thereby exposing the entrapped NBD-glucose to dithionite, and the fluorescent signal fell to zero (**Fig. 3b**, 500 sec time point). This fluorescent signal corresponds to the entrapped NBD-glucose in the nonleaky liposomes.

**Figure 3.**
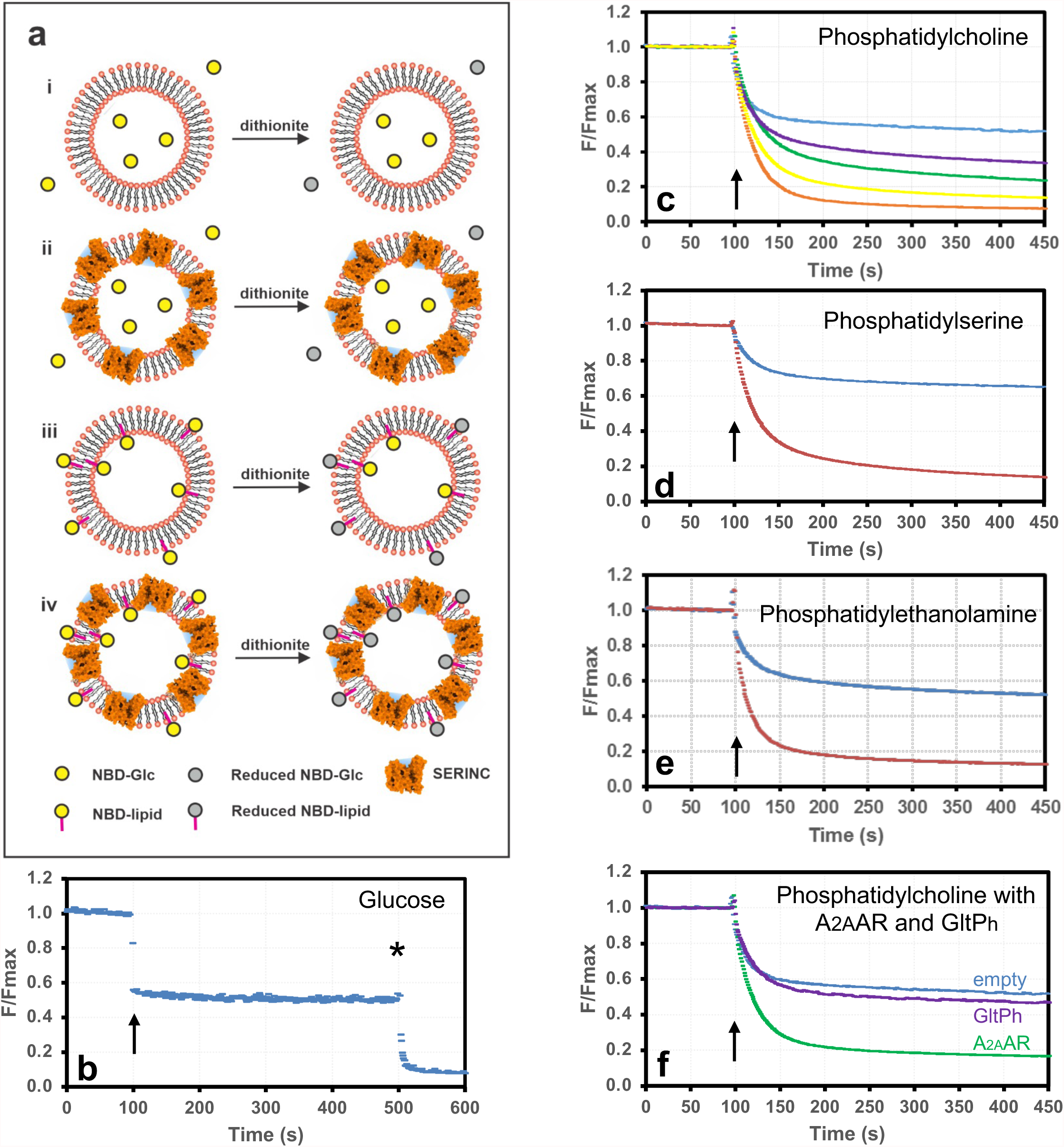
Fluorescent proteoliposome assay demonstrates that hSERINC3 exhibits lipid flipping activity for PC, PE and PS. **(a)** Cartoon showing that the membrane-impermeable, reducing agent dithionite eliminates NBD fluorescence. hSERINC is displayed as being inserted randomly inside-out/outside-in, with blue showing the water cavity between the *α*-helical bundles. NBD-glucose assay shows that (i) empty liposomes and (ii) hSERINC3-containing liposomes are not leaky. (iii) Exterior leaflet NBD-lipids should be accessible to dithionite resulting in ∼50% reduction in fluorescence. (iv) Liposomes containing hSERINC3 should expose inner leaflet NBD- lipids for reduction by dithionite, resulting in ∼100% loss of fluorescence. **(b)** NBD glucose assay demonstrates that liposomes containing a high concentration of hSERINC3 (1.5 µg/mg lipid) are not leaky. The dithionite reduces the fluorescence of extravesicular NBD glucose (60 µM). The stable fluorescent signal from 100 to 500 sec indicates protection of the encapsulated NBD-glucose from dithionite. Confirmation of entrapment was indicated by the reduction of fluorescence to near zero upon solubilization of the liposomes at 500 sec by addition of Triton X-100 (indicated by *). (**c**) - (**f**) Arrows indicate addition of dithionite at 100 sec. The blue curves correspond to the fluorescent profiles for empty liposomes, which display a 40-50% loss of fluorescence upon addition of dithionite. (**c**) Reduction in the fluorescent signal is directly related to the hSERINC3 concentration in liposomes containing NBD-PC. Blue, purple, green, yellow and orange curves correspond to 0.0, 0.25, 0.5, 1.0 and 1.5 µg/mg lipid of hSERINC3, respectively (assuming 100% reconstitution of protein). (**d**) and (**e**) The fluorescent signals of NBD-PE and NBD-PS, respectively, drop to near-zero upon addition of dithionite to liposomes containing 1.5 µg/mg lipid of hSERINC3. (**f**) The adenosine receptor A_2A_AR (1.5 µg/mg lipid) is a known lipid transporter and serves as a positive control for the assay (green). The glutamate transfer homolog GltpH (1.5 µg/mg lipid) has previously been shown to not have lipid flipping activity^29^ and serves as a negative control for the assay (purple). (**b**)-(**f**) Data are representative fluorescence traces of at least 3 independent experiments.

For assessment of lipid flipping activity, proteoliposomes are generated with a fluorescent lipid (NBD-PS, NBD-PC or NBD-PE), which is dispersed randomly in the inner and outer bilayer leaflets amongst the bulk lipids. For empty liposomes, the fluorescence signal should decrease to ∼0.5 since the outer leaflet lipids will be reduced by membrane impermeable dithionite, whereas the inner leaflet lipids will be protected from exposure to dithionite (**Fig. 3c-f**, blue curves). The stable fluorescence from 350-450 sec demonstrates that the empty liposomes were not leaky, in which case the fluorescent signal would have been substantially < 0.5. In the presence of a lipid transporter, one would expect the fluorescent signal to be reduced to near 0 since the inner leaflet lipids will be flipped to the outer leaflet where they are accessible for reduction by dithionite. In our case we used the A_2A_ adenosine receptor (A_2A_AR) as a positive control, which is a known lipid transporter^28^ (**Fig. 3f**, green curve). Our negative protein control was GltpH ^29^, which displayed a fluorescence decay curve (**Fig. 3f**, purple curve) that was close to that of empty liposomes (**Fig. 3f**, blue curve).

In the presence of hSERINC3, the fluorescent signal of NBD-PC, NBD-PE and NBD-PS was reduced to ∼10%, which demonstrates that hSERINC3 manifests lipid flipping activity for PC, PE and PS (**Fig. 3c, d** and **e**, orange curves, respectively). The reduction in fluorescence was related directly to the concentration of hSERINC3 included during liposome reconstitution (**Fig. 3c**). Thus, we conclude that hSERINC3 is a non-ATP-dependent, nonspecific lipid transporter for PC, PE and PS. We suspect that hSERINC3 flips lipids in both directions and functions as a scramblase.

## HIV-1 and MLV particles containing antiviral hSERINCs display elevated levels of phosphatidylserine, which is counteracted by Nef and GlycoGag, respectively

hSERINC5 and to a lesser extent hSERINC3 reduce the infectivity of HIV-1 particles lacking Nef (**Fig. 4a**). In contrast, hSERINC2 lacks antiviral activity. In addition, hSERINC5 mutations have been described that are efficiently expressed and incorporated into viral particles; however, they display varying degrees of impaired antiviral activity^16^ (**Fig. 4a, Extended Data Fig. 9a)**.

**Figure 4.**
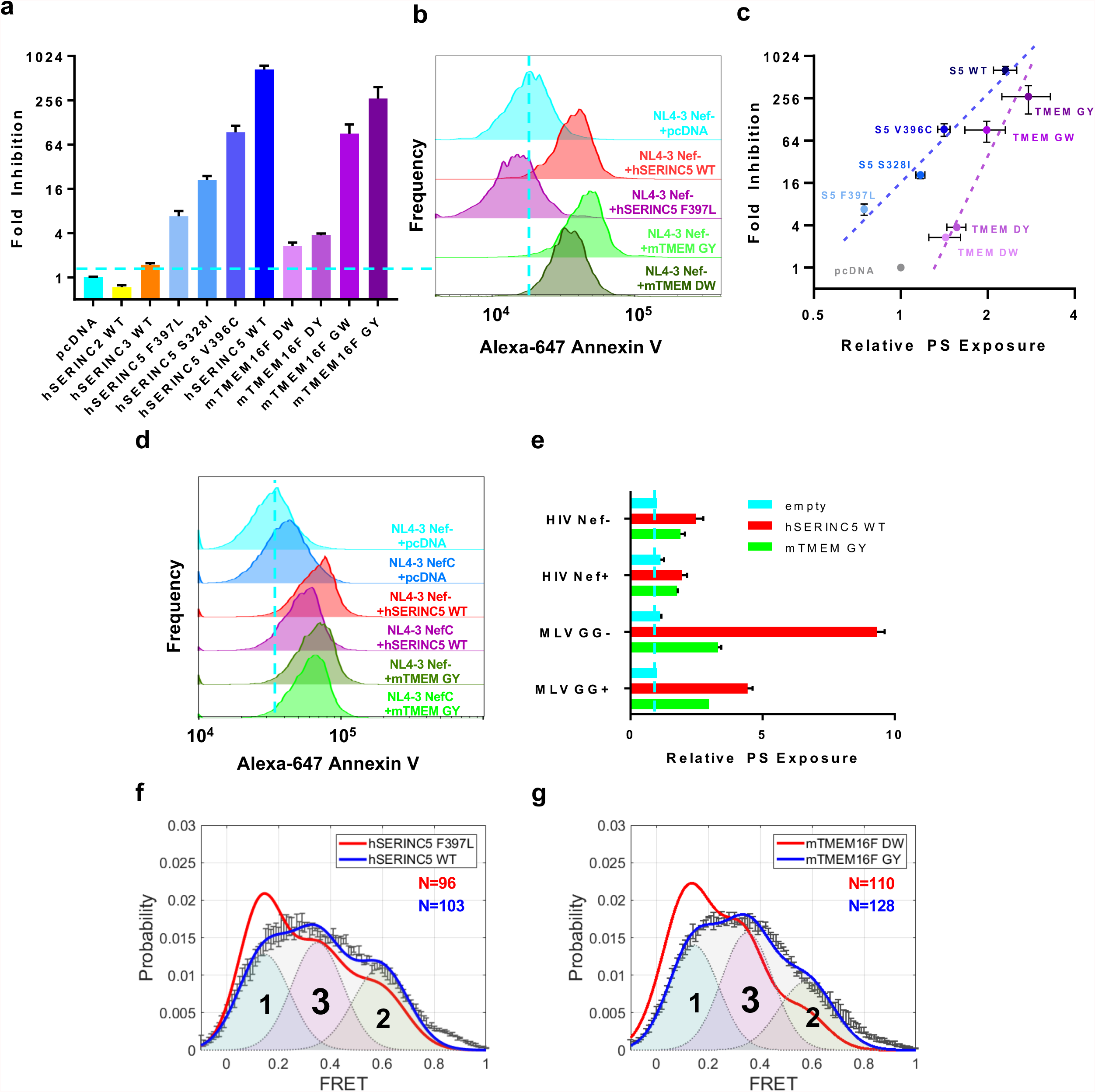
PS exposure by hSERINC5 on virus particles correlates with inhibition of infectivity and conformational changes in Env, and is antagonized by HIV-1 Nef and MLV GlycoGag. **(a)** HIV-1_NL4-3_*Δ*Nef particles were produced in the presence of the indicated plasmid and titered on TZM-Bl indicator cells by Gaussia luciferase assay. Results represent the mean of 7 independent experiments ± SEM. **(b)** HIV-1_NL4-3_*Δ*RT*Δ*Nef virus particles containing Gag-GFP and CD63-mRFP were produced in the presence of the indicated hSERINC5 or mTMEM16F plasmid, bound to anti-CD63 magnetic beads, and stained with Alexa647-annexin V. Virus-bead conjugates were imaged by flow cytometry. Histograms depict annexin V intensity for the GFP- positive population. Results reflect one representative experiment of 3 biological replicates. (c) PS exposure was assessed as in (b) and plotted in relation to virus infectivity as in (a), measured by Gaussia luciferase assay in TZM-Bl cells, for the indicated hSERINC5 (S5) or mTMEM16F proteins (TMEM GY – fully active, TMEM GW – partially active, TMEM DY and DW – inactive). Values represent mean ± SD from 3 independent experiments. (d) HIV-1_NL4-3_*Δ*Nef or NefC- expressing virus particles containing GFP-Vpr and CD63-mRFP were produced in the presence of the indicated hSERINC5 or mTMEM16F plasmid, stained with Alexa647-annexin V, and fixed with 4% PFA prior to analysis by flow cytometry. Histograms depict annexin V intensity for the GFP-positive population from one representative of 3 independent experiments. (e) Annexin V mean fluorescence intensity is shown for HIV-1 particles ± Nef measured by flow cytometry as in (d). Values represent the mean ± SEM from 3 independent experiments. Percent annexin V- positive MLV particles imaged by confocal microscopy as in (Extended Data Fig. 8). Values represent the mean ± SEM from 5 independent experiments. (f) HIV-1_NL4-3_*Δ*RT*Δ*Nef virus particles were produced in the presence of WT hSERINC5 (active, blue) or the F397L (inactive, red) mutant and the conformational state of HIV-1 Env was analyzed by single-molecule FRET (see also Extended Data Fig. 9). N is the number of individual dynamic molecule traces complied into a population FRET histogram (gray lines) and fitted into a 3-state Gaussian distribution (solid) centered at ∼0.15-FRET (dashed cyan), ∼0.35-FRET (dashed red), and ∼0.6-FRET (dashed green). (g) HIV-1_NL4-3_*Δ*RT*Δ*Nef virus particles were produced in the presence of the mTMEM16F GY (active) mutant or mTMEM16F DW (inactive) mutant and the conformational state of HIV-1 Env was analyzed by single-molecule FRET as in (f).

PS is asymmetrically distributed across plasma membranes, with a greater fraction localized to the inner bilayer leaflet. Loss of PS asymmetry plays an important role in a number of biological processes, including apoptosis, blood coagulation and bone mineralization^30^. For our purposes, we used exposure of PS on viral particles as a read-out of lipid flipping, which was detected using annexin V antibodies. Since spontaneous flipping of PS in biological membranes is nearly zero, an increase of exposed PS would indicate the activity of lipid transporters^19, 25^. We generated HIV-1_NL4-3_*Δ*Nef virus particles in the absence or presence of hSERINC5. Incorporation of Gag-GFP was used to label virus particles. We also co-transfected the tetraspanin CD63-mRFP, which was efficiently incorporated and enabled immuno-isolation of virus particles. Two days after transfection, virus particles were harvested from the culture supernatant, bound to anti-CD63 magnetic beads and stained with Alexa647-annexin V. Virus-bead conjugates were monitored by flow cytometry (**Fig. 4b**). The incorporation of hSERINC5 enhanced the exposure of PS on the surface of virus particles (**Fig. 4b,c**). The mutant hSERINC5 F397L exhibited the lowest antiviral activity (**Fig. 4a**) and failed to expose PS (**Fig. 4b,c**). Testing additional hSERINC5 mutants with varying degrees of antiviral effects resulted in a direct correlation between the antiviral activity and the degree of PS exposure on the surface of virus particles (**Fig. 4c**).

To corroborate these findings, we used a second assay to assess PS flipping. HIV-1 particles were labeled by co-transfection of GFP-Vpr, which binds to the p6 domain of Gag and is efficiently incorporated into HIV particles. The exposure of PS was visualized microscopically by Alexa647-annexin V binding to GFP-positive viral particles immobilized on polylysine coated coverslips. hSERINC3 and hSERINC5 enhanced PS exposure, while hSERINC2 did not (**Extended Data Fig. 7a,d**). To confirm that this result was not caused by overexpression of exogenous hSERINC proteins, we compared Gag-GFP labeled particles produced in either parental Jurkat TAg cells or hSERINC3/5 knockout Jurkat TAg cells ^3, 10, 31^. Virus particles produced in parental Jurkat cells showed significantly higher levels of PS exposure than those produced in hSERINC3/5 knockout cells (**Extended Data Figure 7b,e**).

The retroviral accessory proteins Nef and GlycoGag prevent the incorporation of SERINCs into HIV and MLV, respectively, and rescue infectivity^2, 3, 5, 6^. Correspondingly, the exposure of PS on virus particles was abrogated when HIV-1_NL4-3_ expressed the Nef gene from a clade C isolate of HIV-1 (**Fig. 4d**). Similarly, PS exposure was increased in xenotropic MLV, which incorporates an envelope glycoprotein (Env) that is sensitive to SERINCs (**Fig. 4e**). PS exposure was largely reversed when the MLV expressed GlycoGag (**Fig. 4e, Extended Data Fig. 8a,c**). Collectively, these data demonstrate that in two retroviruses, HIV-1 and MLV, SERINC incorporation reduces infectivity and exposes PS, which is counteracted by the retroviral accessory proteins Nef and GlycoGag, respectively.

## An active mutant of the phospholipid scramblase TMEM16F exposes PS on the surface of HIV-1 and MLV and reduces virus infectivity

To fortify the concept that lipid flipping on particles correlates with antiviral effects, we examined the effects of an unrelated PS scramblase, murine transmembrane protein 16F (TMEM16F)^32^. TMEM16F is tightly regulated and only activated during cellular processes such as apoptosis by elevated intracellular calcium levels^33^. During HIV-1 infection TMEM16F in target cells is activated through its calcium-regulated pathway to facilitate membrane fusion^34^. To investigate a possible detrimental role of PS exposure on the surface of the virus, we utilized a constitutively active murine variant (mTMEM16F)^32, 35^, which contains an insertion of 21 amino acids in the amino-tail and a D430G point mutation (designated mTMEM16F GY). The mutant overcomes calcium regulation, displays enhanced PS scrambling activity and is constitutively active^35^. Similar to SERINCs 3 and 5, HIV-1 and MLV containing mTMEM16F GY displayed reduced infectivity (**Fig. 4a,c, Extended Data Fig. 8d** and increased PS exposure (**Fig. 4b,c, Extended Data Figs. 7c,f, 8a-c**). We also assessed the functional consequences of partially active or inactive mTMEM16F mutants (see Methods for details)^36^. In contrast to the fully active mTMEM16F GY mutant, the mTMEM16F GW mutant displayed modestly reduced levels of PS exposure and infectivity inhibition (i.e., partially active), while the mTMEM16F DW and DY mutants displayed severely reduced levels of PS exposure and infectivity inhibition (i.e., inactive). Taken together, these results demonstrate a correlation between PS exposure activity and infectivity inhibition for both hSERINC5 and mTMEM16F variants (**Fig. 4a-c**). As expected, the retroviral accessory proteins Nef and GlycoGag failed to efficiently counteract the effects of mTMEM16F GY (**Fig. 4d,e, Extended Data Fig. 8c, d**). This also concurs with the observation that wild-type TMEM16F is not active in the plasma membrane under normal physiological conditions and is not incorporated into virus particles (**Extended Data Fig. 9a**), and is thus not evaded by retroviruses. The finding that mTMEM16F GY remained active against ecotropic MLV, which evolved to be resistant to hSERINC5, could be related to the inability of viruses to have evaded TMEM16F (**Extended Data Fig. 8c,d**). Importantly, expression of SERINCs and TMEM16F was not toxic to cells, and the absence of effects on cell viability is in agreement with previous results^35^ (**Extended Data Fig. 9b**).

## hSERINC5 and mTMEM16F GY elicit changes in the conformation of the HIV-1 Env trimer

Incorporation of hSERINC5 into HIV-1 particles exposes CD4-induced epitopes on Env glycoprotein trimers such as the membrane proximal external region (MPER) ^31^. We used (1) single molecule FRET (smFRET) and (2) a virus capture assay to explore this observation. smFRET has revealed that HIV-1 Env resides in a prefusion state (designated State 1), which opens in response to CD4 through a necessary intermediate (State 2), into the CD4-bound, open conformation (State 3)^37, 38^. The virus capture assay gives a readout for the binding of conformation-specific antibodies^39^. Incorporation of hSERINC5 resulted in a redistribution of the conformational landscape from State 1 to the State 2 and State 3 conformations (**Fig. 4f, Extended Data Fig. 9f,j**). hSERINC3 exhibited only a slight change in the conformational landscape, and hSERINC2 exhibited no effect (**Extended Data Fig. 9d,e,j**). Correspondingly, the virus capture assay showed that hSERINC3 and, to an even greater extent, hSERINC5 increased access to CD4- induced epitope recognized by 19b (**Extended Data Fig. 9k**). The exposure of CD4-induced epitope recognized by 17b was observed for SERINC5, but not the F397L mutant (**Extended Data Fig. 9k**). The gradual increase in the exposure of antibodies 17b and 19b epitopes in response to incorporation of hSERINC5 mutants correlated with their ability to expose PS and inhibit HIV-1 (**Fig. 4c, Extended Data Fig. 9k**). Moreover, mTMEM16F GY, but not mTMEM16F DW similarly exhibited a profound effect on Env conformation (**Fig. 4g, Extended Data Fig. 9h-j**). These data illustrate that antiviral SERINCs and TMEM16F proteins also affect the conformation of the HIV-1 Env trimer, which presumably occurs as a result of a disturbance in the lipid bilayer associated with lipid transport.

## Purified SERINC5 and SERINC2, as well as SERINC5 mutants F397L and S328I also flip lipids

Unlike the action of hSERINC5 on virus particles, neither hSERINC5 mutants F397L and S328I, shown in **Extended Data Fig. 6i,j**) nor hSERINC2 enhance the exposure of PS, affect Env conformation or reduce infectivity. We also used the proteoliposome assay to assess the lipid flipping activity of wild type hSERINC5, the mutants F397L and S328I, as well as hSERINC2. Interestingly, all proteins exhibited similar lipid flipping activities (**Extended Data Fig. 2l**). Although the hSERINCs exhibited similar lipid flipping, fluorescence decay curves (**Figure 3c-e**, **Extended Data Fig. 2l**), quantitative analysis (**Extended Data Fig. 2m**) showed that there are differences in the flipping rates attributable to the fast component of dithionite reduction: hSERINC3 flips fastest (5.84 ± 0.15) × 10^-2^ s^-1^), hSERINC5 flips the slowest (1.92 ± 0.06) × 10^-2^ s^-1^), and hSERINC2 flips at an intermediate rate (2.87 ± 0.08) × 10^-2^ s^-1^).

We realize the discordance between the preserved lipid flipping activity in proteoliposomes containing hSERINC2 or the hSERINC5 mutants (F397L and S328I) and the reduced lipid flipping and reduced restriction in HIV-1 particles containing hSERINC2 and the hSERINC5 mutants. The appeal of the proteoliposome assay is the elegant simplicity and chemical definition of the system.

The liposomes are formed from synthetic lipids (POPC and POPG) doped with a fluorescent lipid, and the proteoliposome membrane contains a single, purified protein species. However, the proteoliposomes do not recapitulate several features present in cell membranes and the HIV-1 envelope. For instance, the lipids are symmetrically distributed upon generation of proteoliposomes, whereas membranes display increased concentration of PS, PE, PIPs and sphingomyelin in the cytoplasmic leaflet, and glycolipids are concentrated in the extracellular leaflet^40, 41^. In addition, PIP2 and PIP3 are enriched in the HIV-1 envelope compared with the plasma membrane (Mücksch *et al.*, Quantification of phosphoinositides reveals strong enrichment of PIP2 in HIV-1 compared to producer cell membranes. *Sci. Reports* 9:17661 (2019) PMID 31776383). The transporters (e.g., hSERINCs and TMEM16F) are symmetrically distributed in their orientation upon reconstitution, whereas the asymmetric orientation of the proteins in the plasma membrane is retained upon particle budding. In addition, HIV-1 Env and SERINCs reside in microdomains^31, 42^, which are not present in proteoliposomes. Other proteins such as the tetraspanin CD63 reside in microdomains within the envelope^43^. These distinguishing features in the HIV-1 envelope may contribute to the restriction phenotype, not present in proteoliposomes.

## A hypothetical model for SERINC-mediated lipid flipping

Given the lipid flipping activity of hSERINC3, we performed MD simulations and AlphaFold analysis to explore the conformational dynamics, lipid interactions and potential mechanism of action. Unlike proteoliposomes, the composition of the lipid bilayers was designed to be similar the HIV-1 fusion environment^44^. The lipid bilayer was generated with an equal distribution of lipids in each leaflet, except for PIP2 and sphingomyelin, which were restricted to the cytoplasmic and extracellular leaflets, respectively. Following the equilibration phase, it was reassuring that there were stable binding sites for lipids (**Fig. 5b,c,d**), and the tertiary structures of the helical bundles and the H4 crossmember helix remained stable (**Fig. 5a-e**), with backbone RMSD values of 2.0 – 2.5 Å. A notable structural change during equilibration was significant thinning of the bilayer from ∼45 to ∼30 Å (**Fig. 5a**). Bilayer thinning has been observed for other membrane transporters^45, 46^ and has been shown to be critical for lipid flipping of TMEM16f^47, 48^.

**Figure 5.**
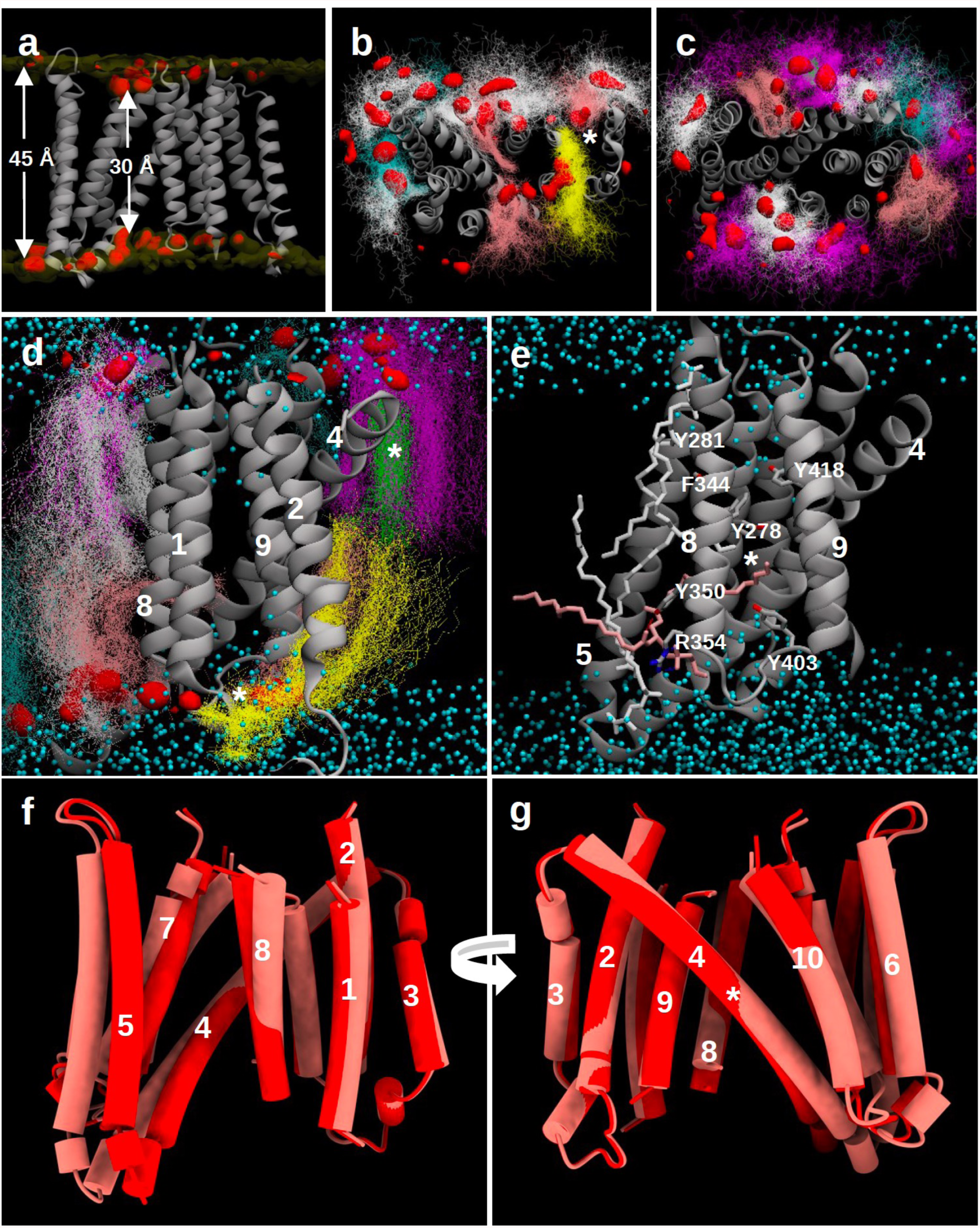
MD simulations and AlphaFold analysis of hSERINC3 display bilayer thinning, lipid and cholesterol binding sites, a water channel and conformational changes suggesting an alternating access mechanism. Four simulations were performed for a total of 4.1 µs, starting with different random starting positions of the lipids. <u>Bilayer thinning</u>: (a) Shown is the density of lipid phosphorus atoms during a 2.1 µs hSERINC3 (gray ribbon) simulation with the highest density red and an intermediate density transparent gold. The lipid bilayer thins from ∼45 to ∼30 Å, with the narrowest constriction across the H5/H8 cleft. <u>Lipid and cholesterol binding</u>: Cytoplasmic (b), extracellular (c) and side views (d) of hSERINC3. Thin lines show the positions of lipid molecules at 20 ns intervals (POPS, white; POPE, pink; POPC, cyan; sphingomyelin, magenta; PIP2, yellow; cholesterol, green). A boundary layer of lipid and cholesterol molecules displayed stable interactions with the hydrophobic surface of hSERINC3 (b-d) including hydrogen bonding interactions between PIP2 molecules and basic residues at the cytoplasmic periphery of the protein (b,d, yellow). In some cases lipids interact with the protein via bound cholesterol molecules (e.g., asterisk in (d) showing sphingomyelin/cholesterol interactions). <u>Water channel</u>: (d,e) Side views show a water channel between H4 and H8 that can span the bilayer, which may be a putative transport pathway. The entrances to the water channel correspond to the narrowest constriction of the bilayer. (e) The water channel is bounded by a series of tyrosine residues (Y278, Y281, Y350, Y403, Y418) and a phenylalanine residue (F344), some of which are involved in stable lipid binding. For instance, Y350 and R354 form hydrogen bonds with a POPE molecule during most of the 2.1 µs simulation. The asterisk in (e) identifies the location of a satellite density visible in Fig. 2c, consistent with a bound lipid, cholesterol or detergent molecule., and may indicate a possible transport pathway. <u>Conformational changes suggesting an alternating access mechanism</u>: (f,g) The top two conformational states of hSERINC5 determined by AlphaFold displayed motions between the helical bundles accompanied by a hinge-like movement of the H4 crossmember helix consistent with an alternating access mechanism. Also see Extended Data Movies 1 and 2.

MD simulations of MurJ^23, 49^ and TMEM16f^47, 50^ displayed deep penetration of water into the bilayer. Likewise, during our simulations of hSERINC3, water molecules penetrated the interface between the crossmember helix H4 and its companion helix H8 (**Fig 5d,e**), which was also observed in MD simulations of TMS1d^16^. Notably, in our case, there were instances during which a continuous chain of water molecules traversed the hydrophobic zone of the bilayer **(Fig. 5d)**, which may correspond to a transport pathway. Interestingly, the aqueous pathway in hSERINC3 is bounded by a series of tyrosine residues (Y278, Y281, Y350, Y403, Y418) and a phenylalanine residue (F344), some of which are involved in lipid binding (**Fig. 5e**). For instance, Y350 and R354 form hydrogen bonds with a POPE molecule during most of a 2.1 µs simulation. The cryoEM map also displayed a satellite density adjacent to aromatic residues F182, W186 and W190 at the cytoplasmic ends of H4 and H5 **(Fig. 2c)**. The shape of the density was consistent with a bound molecule of lipid, cholesterol or detergent, which may be a putative transport pathway. Cholesterol is known to associate with lipid acyl chains^51^, and we observed instances during the simulations in which cholesterol molecules bound directly to the protein or interacted stably with lipids bound to hSERINC3 (e.g., cholesterol (green) * in **Fig. 5d** and sphingomyelin (magenta)).

For MurJ, lipid II is proposed to enter the protein via a lateral gate, and the crossmember *α*-helix 7 serves as a fulcrum that may facilitate flipping of lipid II to the outer leaflet (**Extended Data Fig. 1i,j**)^23^. The cryoEM maps of hSERINC3 show that the helical bundles may move as rigid bodies (**Fig. 1f**) with a pivot point in the center of the crossmember helix H4, which were also observed in AlphaFold analysis of hSERINC3 (**Fig. 5i,j, Extended Data Movies 1,2**). These movements are akin to the lever-arm-like movement of helix 7 in the MurJ flippase **(Fig. 5i,j)**^23^.

For PfMate ^21^ and MurJ ^52^, lipid flipping has been proposed to involve a mechanism similar to alternating access of membrane transporters. Such a mechanism has also been suggested for LtaA ^24^ but without experimental evidence. For MurJ, an alternating access mechanism is supported by multiple X-ray structures^23^ and cross-linking experiments^52^. However, MD simulations of MurJ did not show flipping of lipid II. Likewise, we did not observe lipid flipping during our simulations of hSERINC3. This is not surprising given the slow time scale of lipid flipping compared with the microsecond time scales of MD simulations^18, 20^. Nevertheless, the top two conformational states of hSERINC5 predicted by AlphaFold displayed motions between the helical bundles accompanied by a hinge-like movement of the H4 crossmember helix consistent with an alternating access mechanism **(Fig. 5i,j)** and **Extended Data Movies 1** and **2**.

## Conclusions

In summary, the evidence supporting the hypothesis that SERINC-mediated lipid flipping is relevant to the mechanism of restriction include the following: (1) The architecture of hSERINC3 **(Figs. 1,2)** resembles that of other non-ATP dependent lipid transporters such as PfMATE^21^, MurJ^22, 23^ and LtaA^24^. (2) Our comparison of the full-length and *Δ*ICL4-constructs of hSERINC3 shows rigid-body rotation of the helical domains and bending of the H4 “crossmember” helix (**Fig. 1f**). These features are similar to the conformational changes of MurJ in the flipping of Lipid II (**Extended Data Fig. 1i,j**)^23^. (3) MD simulations displayed water penetration through the hydrophobic zone and binding sites for PS, PC, PE and cholesterol, as well as thinning of the bilayer, which has been shown to be critical for the lipid flipping activity of TMEM16^53^ (**Fig. 5a- e**). (4) Purified hSERINC3 incorporated into proteoliposomes exhibits flipping activity for PC, PE and PS (**Fig. 3 c,d,e**). Nonspecific lipid flipping is a hallmark of non-ATP dependent lipid scramblases. (5) Annexin binding to HIV-1 and MLV particles assembled in the presence of hSERINCs3 and 5 (but not 2), indicates that PS, which is normally enriched on the inner bilayer leaflet, is exposed on the outer leaflet of the viral membrane (**Fig 4b,c, Extended Data Figs. 7a,7d, 8a-c**). (6) Virus particles produced in parental Jurkat cells showed significantly higher levels of PS exposure than those produced in hSERINC3/5 knockout cells (**Extended Data Fig. 7b,e**). (7) The accessory proteins Nef and GlycoGag, known to prevent SERINC from being incorporated into viral particles, reduce PS levels on the virus surface (**Fig. 4c,d, Extended Data Fig. 8a,c**). (8) SERINC5 mutants with impaired antiviral activity display reduced PS exposure (**Fig. 4a-c**). (9) The constitutively-active mTMEM16F GY mutant is incorporated into viral particles, exposes PS on virus particles and reduces virus infectivity comparable to that of SERINC5 (**Fig. 4b,c**) (10) SERINC and TMEM16F proteins with antiviral activity alter the conformation of Env (**Fig, 4f,g, Extended Data Fig. 9f,i-k**).

Our results using two lipid flipping proteins (hSERINCs and mTMEM16f) and two retroviruses (HIV-1 and MLV) show that lipid flipping is directly correlated with changes in Env conformation and restriction activity. Thus, we conclude that SERINCs are nonspecific, non-ATP-dependent, phospholipid transporters, and we hypothesize that lipid flipping elicits antiviral effects that culminate in reduced infectivity.

## Acknowledgments

We thank Shigekazu Nagata for the mTMEM16F plasmids, Felipe Diaz-Griffero for hSERINC2 plasmid, Heinrich Göttlinger for Jurkat TAg and SERINC3/5 knockout cells, and Tom Hope for the GFP-Vpr plasmid. We thank Olga Boudker (Weill Cornell) for providing purified GltpH. CryoEM data were primarily collected at the University of Virginia Molecular Electron Microscopy Core facility (RRID:SCR_019031), which was established via recruitments funds to M.Y., NIH grant G20 RR31199, NIH grant S10-RR025067 and NIH grant S10-OD018149. CryoEM data were also collected with the assistance of William Rice at the National Center for CryoEM Access and Training (NCCAT) and the Simons Electron Microscopy Center located at the New York Structural Biology Center, which is supported by the NIH Common Fund Transformative High Resolution Cryo-Electron Microscopy program (U24 GM129539), and by grants from the Simons Foundation (SF349247) and NY State.

## Data and materials availability

The atomic coordinates and EM maps for WT hSERINC3-Fab and *Δ*ICL4-hSERINC3-Fab have been deposited in the Protein Data Bank (www.rcsb.org) and EMDB (www.ebi.ac.uk/pdbe/emdb/) with accession codes 7RU6 and EMD-24698 and 7RUG EMD-24705, respectively.

## Contributions

S.A.L., S.P. and W.E.M. performed protein expression and purification. S.A.L., S.E., K.N. and A.A.K. generated synthetic Fab proteins. M.D.P. and S.A.L. performed cryoEM, image analysis and model building. M.D.P. performed MD simulations and normal mode analysis. S.A.L. performed the proteoliposome fluorescent lipid flipping experiments with input from M.Y. E.T. performed “curve stripping” analysis of the fluorescence decay curves, with input by M.Y. and S.A.L. J.R.G., Z.Y., S.D., P.D.U., A.F. and W.M. performed virus imaging experiments, flow cytometry, HIV-1 capture and infectivity experiments. M.L., J.R.G. and W.M. performed smFRET studies. K.K.L. and A.R. performed PS exposure experiments in MLV particles. The manuscript was written primarily by S.A.L., M.D.P., J.R.G., W.M. and M.Y., with input from all authors throughout experimentation and manuscript preparation.

**Online Content Methods, along with any additional Extended Data display items and Source Data, are available in the online version of the paper.**

## Methods

### Constructs and expression of human SERINC proteins

#### SERINC3

The gene that encodes human SERINC3 (Genscript-OHu02717D) was inserted upstream of a thrombin protease cleavable linker (LVPRGS) and Strep II epitope (WSHPQFEK) in a modified pFastBac vector by In-Fusion cloning (Clontech). Mutagenesis using the QuikChange Site-Directed Mutagenesis kit (Agilent) was performed to change the thrombin site to a Tobacco Etch Virus (TEV) protease site (ENLYFQ\S) to facilitate expression and purification. QuikChange mutagenesis was also used to remove hSERINC3 amino acids 366-391 from this vector to generate ΔICL4-hSERINC3. For synthetic Fab production, the original hSERINC3 pFastBac plasmid had the thrombin protease linker and Strep II epitope deleted, and a Flag epitope inserted (DYKDDDDK) by site-directed mutagenesis. These constructs were expressed in Sf9 insect cells using the Bac to Bac Baculovirus system (Invitrogen). The cells were infected with baculovirus at 27 °C for 48 hrs before harvesting.

#### SERINC5

The construct design, expression and purification of human SERINC5 recapitulated that of SERINC3 as they were done simultaneously. The gene that encodes human SERINC5 (OHu11910D) was purchased from GenScript.

#### SERINC2

Isoform 1 of human SERINC2 (GenScript-OHu23082D) was cloned into pFastBacI with the TEV and STREP cleavage and affinity tags upstream of the gene encoding hSERINC2. For this purpose we used the SERINC3 construct, in which the SERINC3 open reading frame was removed by restriction enzyme digestion. The hSERINC2 gene was then inserted via ligation after PCR amplification to insert the appropriate restriction site on the 5’ and 3’ ends.

### Formation of hSERINC3 nanodiscs for generation of Fabs

Sf9 cell pellets infected with virus containing C-terminally FLAG-tagged hSERINC3 were lysed in 50 mM HEPES, pH 7.5, 50 mM NaCl, and 0.5 mM EDTA and protease inhibitors (cOmplete Ultra Tablet (Roche)). After lysis, the mixture was incubated with 2.5 mM MgCl_2_ and benzonase (0.5 µl/ml) (EMD Millipore, Corp) for 10 min before centrifugation at 125,000 x *g* for 40 min to collect the membranes. The membranes were washed twice by homogenization in 50 mM HEPES, pH 7.5, 1 M NaCl and 0.5 mM EDTA and were then collected by centrifugation at 125,000 x *g* for 40 min. Membranes were suspended in 40% glycerol, 10 mM HEPES, pH 7.5, 20 mM KCl, and 10 mM MgCl_2_ before flash-freezing in liquid nitrogen and storage at -80 °C for further use.

hSERINC3 was extracted from purified membranes using 1% n-dodecyl-β-D-maltoside (DDM) (Anatrace), 0.2% (w/v) cholesteryl hemisuccinate (CHS) (Anatrace) in the presence of 2.0 mg/ml iodoacetamide and purified by M2 anti-Flag (Sigma) immunoaffinity chromatography. After washing with high salt buffer (1M NaCl) and progressively lowering the DDM/CHS concentration, hSERINC3 was eluted in a buffer consisting of 50 mM HEPES pH 7.5, 150 mM NaCl, 0.02% DDM, 0.004% CHS and 0.2 mg/ml Flag peptide (Bio Basic, USA). β-mercaptoethanol (βME) was added to 2.0 mM before the sample was concentrated using a Vivaspin 50 kDa molecular-weight cut-off filter (GE Healthcare.) The monomeric fractions were purified by size exclusion chromatography using a Superose 6 Increase column in 50 mM HEPES pH 7.5, 150 mM NaCl, 0.02% DDM, 0.004% CHS and 2 mM βME. Before incorporation into nanodiscs, a PD-10 column (GE Healthcare) was used to remove the βME, and the hSERINC3 was concentrated to ∼42 µM. Purified hSERINC3, biotinylated membrane scaffold protein MSP1D1 (provided by the Kossiakoff lab), and soybean polar lipid extract (Avanti Polar Lipids) were mixed at a molar ratio of 1.0:2.5:300. After 1 hr on ice, the sample was subjected to detergent removal with Bio-Beads SM-2 (BioRad). Bio-beads were removed, and the reconstitution mixture was cleared by centrifugation prior to injection onto a Superose 6 Increase column for removal of empty nanodiscs. Peak fractions were evaluated by SDS-PAGE and silver-staining (4-20% Mini-Gel; BioRad) to confirm the presence of both hSERINC3 and MSP1D1. Peak fractions were pooled, adjusted to 5% with glycerol, concentrated and flash frozen for storage at -80 °C until use.

### Phage Display

hSERINC3 was reconstituted into nanodiscs using biotinylated MSP1D1, which was chemically biotinylated and assessed for pull-down efficiency as previously described^1–3^. Five rounds of biopanning were performed using Fab Library E^4, 5^ in a buffer containing 25 mM HEPES, pH 7.4, 150 mM NaCl with 1% BSA (Selection Buffer) using a method adapted from published protocols^1, 2^. In the first round, biopanning was performed manually using 200 nM of hSERINC3- MSP1D1 nanodiscs immobilized onto Streptavidin paramagnetic beads (Promega). Following three washes with Selection Buffer, the beads were directly used to infect log-phase *E. coli* XL-1 Blue cells, and the phage pool was amplified overnight as described^2^. To increase selection stringency, four additional rounds of sorting were performed semi-automatically using a Kingfisher magnetic beads handler (Thermo Fisher Scientific), with the amplified phage pool from each preceding round used as input. For each of these rounds, target concentrations were decreased as follows: 2nd round: 100 nM; 3rd round: 50 nM; 4th round: 50 nM; and 5th round: 25 nM. The phage pools for rounds 2-5 were pre-cleared with 100 µL of streptavidin particles, and for all rounds between 1.0 µM and 1.5 µM, empty, non-biotinylated MSP1D1 nanodiscs were used as soluble competitors. The 4th and 5th round phage pools were also tested against immobilized 25 nM empty, biotinylated MSP1D1 nanodiscs and streptavidin beads alone to evaluate hSERINC3- specific enrichment. For rounds 2-5, bound phage particles were removed by elution with 1% Fos-choline-12 as described^1^.

### Single-point Phage ELISA

Single-point phage ELISA was used to evaluate individual clones from rounds 4 and 5. All ELISAs were performed in 96-well plates (Nunc) coated with 2 µg/mL Neutravidin and blocked with Selection Buffer containing 1% BSA. Colonies of *E. coli* XL-1 blue harboring phagemid were used to inoculate 2xYT media supplemented with 100 µg/mL ampicillin and 10^9^ pfu/mL M13-KO7 helper phage. The phage were amplified overnight in deep well blocks at 37 °C with shaking at 280 RPM. Amplified phage were diluted 10-fold into Selection Buffer and assayed against hSERINC3-loaded 1D1 nanodiscs or empty biotinylated nanodiscs at 25 nM concentration. ELISA was performed as described^2^ using an HRP-conjugated anti-M13 monoclonal antibody (GE Healthcare) and a TMB substrate kit (Thermo Fisher) to detect bound phage. Wells containing buffer alone were also used to determine the specificity of binding.

### Fab expression and purification

hSERINC3-specific binders from phage ELISAs were selected based on signal/background ratios ^1^ and sequenced at the University of Chicago Comprehensive Cancer Center DNA Sequencing facility. Unique clones were sub-cloned in pRH2.2 (a gift of S. Sidhu) using the In-Fusion Cloning kit (Takara) and sequence-verified. Fab expression vectors were then transformed into *E. coli* BL21-Gold competent cells (Agilent), and Fabs were expressed as described^1, 2^. Cells were harvested by centrifugation, and the cell paste was frozen until use. Fabs were purified as previously described^1, 2^.

### Purification of hSERINC3 and hSERINC3-Fab complex formation

Strep-tagged hSERINC3 or ΔICL4-hSERINC3 were extracted from purified membranes by incubation for 2-3 hrs at 4 °C in 1% DDM, 0.2% (w/v) CHS in the presence of 2.0 mg/ml iodoacetamide in 50 mM HEPES, pH 7.5, 300 mM NaCl and 2.5% glycerol. The unextracted material was removed by centrifugation at 125,000 x *g* for 40 min. The supernatant was incubated overnight at 4 °C with Strep-Tactin Beads (Qiagen), typically using 0.75 ml packed beads per two liters of original culture volume. After binding, the beads were washed with high salt buffer (1.0 M NaCl) containing a progressively lower DDM/CHS concentration (0.05% DDM/0.001% CHS-final). The beads were then exchanged into high salt buffer (500 mM NaCl) containing 0.5% glyco-diosgenin (GDN, (Anatrace)) and incubated with PNGaseF (New England Biolabs) for 1 hr at room temperature. The GDN concentration was lowered by one wash in high salt buffer (500 mM NaCl) containing 0.1% GDN, and hSERINC3 was eluted from the beads with desthiobiotin (Sigma) in low salt buffer (150 mM NaCl) containing 0.0045% GDN. The desthiobiotin was removed using a G-25 buffer exchange column (GE Lifescience). The hSERINC3 concentration was determined and mixed with a 1.25 molar excess of the Fab on ice for 1 hr before the addition of the same Strep-Tactin beads used previously, but which had subsequently undergone regeneration. The bead mixture was incubated overnight at 4 °C under gentle agitation. The beads were subsequently washed with high salt buffer (1.0 M NaCl), followed by low salt buffer (150 mM NaCl), with both buffers containing 0.0045% GDN. hSERINC3 was eluted in low salt buffer contain 2.5 mM desthiobiotin and then concentrated using a centrifugal filter (Vivaspin, 50 kDa MWCO (GE Healthcare)). The solution was subjected to a 5 min spin at 17,000 x *g* in a refrigerated microcentrifuge before separation on a Superdex 200 10/300 column (GE Healthcare) equilibrated with SEC buffer (50 mM HEPES, pH 7.5, 150 mM NaCl, 0.0045% GDN). Peak fractions were subjected to silver stain analysis to confirm the presence of both hSERINC3 and the Fab. Selected monomer peak fractions with the Fab were pooled and concentrated.

### CryoEM grid preparation and data collection

All grids for cryoEM were prepared in the University of Virginia Molecular Electron Microscopy Core (MEMC). C-Flat 1.2/1.3 400C grids (Protochips) were glow discharged in a Pelco EasiGlow for 45 s at 20 mA. A 2.1 µL aliquot of WT hSERINC3-Fab at ∼4 mg/mL was applied to the carbon side of each grid in a Vitrobot Mark IV held at 4°C and 95 % relative humidity. Grids were blotted in the Vitrobot with Whatman #1 filter paper, using a blot force of 2 and blot time of 7 s. The WT SERINC3 sample was then vitrified in liquid ethane cooled by liquid nitrogen. *Δ*ICL4-SERINC3-Fab was crosslinked with 10 mM BS3 for 45 min at room temperature, concentrated to ∼4 mg/mL, and cryoEM grids were prepared in the same way, with a blot force of 5 and blot time of 7 s. Suitable grids and grid regions of hSERINC3-Fab samples were identified by screening atlas images on the UVa MEMC Titan Krios (Thermo Fisher Scientific) using the Falcon 3EC detector. CryoEM data collection of the best grids was performed at the New York Structural Biology Center (NYSBC) Titan Krios III equipped with a Gatan K2 detector and a GIF energy filter. A priority for cryoEM of hSERINC3- Fab was to record images of the thinnest ice that still contained a suitable number of particles. The optimal ice thickness was 30-40 nm. In ice 25-30 nm thick, particle exclusion was significant, and in ice thinner than 25 nm, all particles were excluded. In ice thicker than 50 nm, the Leginon maximum resolution estimates degraded substantially. Leginon data collection criteria were set to exclude imaging positions with ice thickness less than 25 nm and greater than 60-70 nm using the Leginon Holefinder ice thickness determination method.

All other hSERINC3-Fab and *Δ*ICL4-hSERINC3-Fab cryoEM datasets were collected on the UVa MEMC Titan Krios equipped with a Falcon 3EC detector using EPU automated acquisition software (Thermo Fisher Scientific). Data collection details are summarized in **Extended Data Table 1.**

### CryoEM image processing and reconstruction

All cryoEM image processing and reconstructions were performed using RELION 3.0 or RELION 3.1. An overview of the processing is presented here for the wild type hSERINC3-Fab data collected at NCCAT. Processing was similar for the other datasets, and RELION flowcharts and results for the WT SERINC3-Fab and *Δ*ICL4-hSERINC3-Fab are presented in **Extended Data Figs. 3-5**. Movie frame alignment and dose weighting were performed with MotionCor2 v1.1^6^ using 5 x 5 patches. In the case of the NCCAT data set, images were 2x binned for a final pixel size of 1.298 Å/pixel. Contrast Transfer Function (CTF) estimation was performed with CTFFIND 4.1.13 on the aligned and dose-weighted micrographs. Images that met the following CTFFIND criteria were selected for particle picking: maximum resolution estimates better than 3.8 Å, underfocus range -0.8 – -2.2 µm and astigmatism < 200 Å. Particles from a subset of the data were picked automatically using a Laplacian of Gaussian (LoG) model with minimum and maximum diameters of 120 and 180 Å, respectively. Following extraction, these particles were “cleaned” using 2D classification, the resulting particle set was used to generate an *ab initio* initial model, and then subjected to 3D classification. The best 3D model was used for particle auto-picking from the full dataset. Picked particles (515,230) were extracted with a box size of 200 pixels (259.6 Å) (**Extended Data Fig. 3a)**. The particle set was “cleaned” again using 2D classification with 24 classes. Some 2D classes clearly showed α-helices for hSERINC3, while particles were poorly aligned in other classes. All classes that appeared to include a substantial number of “real” particles were selected for further processing, resulting in 495,552 particles. The same model used for particle picking was low-pass filtered to 15 Å and used for 3D classification with 6 classes and a spherical mask of 180 Å. 3D classification resulted in one good class with 164,700 particles. Particles from this class were subjected to 100 iterations of 3D classification (one class) with the same spherical mask and a regularization parameter of T = 20. The resulting map was low-pass filtered to 15 Å and used to generate a mask with a soft edge of 11 pixels. The mask and the map from 3D classification (low-pass filtered to 4 Å) were used for half-map 3D auto-refinement with an initial angular sampling of 0.9° and an initial offset range of 1 pixel. Aligned particles were subjected to CTF refinement and Bayesian polishing and refined as before with no noticeable improvement in the GSFSC resolution or map appearance.

A significant obstacle to high-resolution refinement of WT hSERINC3-Fab and *Δ*ICL4- hSERINC3-Fab maps was apparent conformational heterogeneity between the two bundles of transmembrane *α*-helices. We tried a variety of masking, particle subtraction, and local refinement strategies and programs (RELION 3.0/3.1^7^, cryoSPARC v2.1^8^, cisTEM 1.0^9^) in attempts to improve the maps. Alignments were dominated by the Fab-proximal helical bundle and the Fab variable domains (designated Fab_v_), which contributed to the reduced resolution of the Fab-distal helical bundle. However, attempts to refine the hSERINC3 helical bundles individually failed due to the small size of each fragment (e.g., Fab-distal helical bundle < 20 kDa). In order to generate the best map for each hSERINC3 helical bundle and measure the approximate resolution of each, we generated separate maps for the proximal bundle-Fab_v_ domain and the distal bundle. The separate maps were used to create a mask for each domain, which were then used for post-processing in RELION. Optimal B-factor sharpening values were determined empirically. B-factors of -75 Å^2^ for the Fab-proximal domain and -50 Å^2^ for the Fab-distal resulted in the best combination of map detail and continuity of density. The resulting maps were combined in Chimera and used for structure analysis and model building.

### Model building and refinement

An initial model was built in Coot 0.8.9.2^10^ using the hSERINC3-Fab_v_ domain map. Ideal *α*-helices were placed in the map using the topology of the Drosophila melanogaster hSERINC homolog TMS1d^11^. The model of the hSERINC3 Fab- proximal bundle (H4-5-6-7-10) was built using bulky side chains and proline densities, combined with MEMSAT-SVN and Protter ^12^ hydropathy and PSIPRED 4.0^13^ secondary structure predictions to determine the sequence register in the map. Putative models of the Fab-distal bundle (H1-2-3-8-9) were built using a similar strategy combined with hydrophobicity and electrostatic analysis of each model in ChimeraX^14^ and analysis of membrane insertion energies with the PPM server^15^.

An initial model of the Fab_v_ domain was generated using the Robetta server ^16^ and the synthetic antibody structure PDB: 5bk1^17^. The Fab_v_ model was manually rebuilt in Coot. The hSERINC3-Fab_v_ model including all side chains was refined against the map in real space using ISOLDE 1.0b3^18^ in ChimeraX. During ISOLDE refinement helical restraints were used on the full length of all transmembrane helices with the exception of H3, which had two discontinuous rods of density and was therefore divided into two helical segments. We then truncated the side chains of the Fab-distal bundle to Cα in the hSERINC3 model. which we determined was most likely to be correct based on agreement with the cryoEM map and energetic and hydrophobicity analysis.

A model of *Δ*ICL4-hSERINC3-Fab_v_ was generated from the hSERINC3-Fab_v_ model by adding an ideal helix to the *Δ*ICL4-hSERINC3 model in the H8 helical density between the two bundles. The H8 helical density had insufficient side chain features to unambiguously position the sequence, so we relied on the MEMSAT-SVM and Protter predictions and energetic and hydrophobic analyses to build and position a probable model for H8. The resulting *Δ*ICL4- hSERINC3-Fab_v_ model was refined in ISOLDE with helical restraints and a low weight on the map contribution to the refinement simulation (10 – 20 x 1000 kJ mol^-1^ (map units)^-1^ Å^3^).

Due to the absence of some side chains in the WT hSERINC3-Fab and most of the side chains in the *Δ*ICL4-hSERINC3-Fab cryoEM maps, we used AlphaFold version 2 (10.1038/s41586-021-03819-2) to generate complete models of the hSERINC3 transmembrane domains. We believe these hybrid cryoEM/AlphaFold models are the best estimations of the hSERINC3 structures. We ran AlphaFold v2.1 on a local workstation with the full database, except for exclusion of the TMS1d structure (PDB:6SP2). We removed low-confidence (low pLDDT scores) loops from the top scoring AlphaFold model, which corresponded to density that was not visualized in the cryoEM maps. All the transmembrane *α*-helices of the AlphaFold model had very-high confidence scores except H8, which was also weaker in the WT cryoEM density map. We docked the top AlphaFold hSERINC3 model into each cryoEM map (WT and *Δ*ICL4) in ChimeraX, then used ISOLDE v1.2 to refine the models. To preserve the AlphaFold side chain information, we used a very low map weight (5 x 1000 kJ mol^-1^ (map units)^-1^ Å^3^), and we used secondary structure restraints for all helical segments. Following ISOLDE refinement, we added the CDR of the bound Fab and performed further refinement of the models using Phenix Refine and real space refinement in COOT. The deposited hybrid models (7RU6 and 7RUG) include all side chains, and details are summarized in **Extended Data Table 1**.

### Purification of hSERINC’s for incorporation into liposomes

Strep-tagged hSERINC3, hSERINC2, hSERINC5, hSERINC5-F397L and hSERINC5-S328I were purified as described above for the hSERINC3-Fab complexes, except that the proteins were not exchanged into GDN. After elution from the Strep-Tactin Beads (Qiagen), the proteins were further purified on a Superdex 200 10/300 column (GE Healthcare) equilibrated with SEC buffer (50 mM HEPES, pH 7.5, 150 mM NaCl, 0.05% DDM/0.001% CHS). Peak fractions were pooled and concentrated to ∼1.0 mg/ml.

### Preparation of liposomes

Preparation of liposomes and proteoliposomes and performance of the lipid flipping assay were adapted from protocols published by the Menon lab^19^. Liposomes and proteoliposomes were prepared in 50 mM HEPES, pH 7.5, and 150 mM NaCl (i.e., liposome buffer). A glass syringe (Hamilton, 50 and 500 µl), was used to add 215.5 µl POPC (25 mg/ml, in chloroform, Avanti) and 24.6 µl POPG (25 mg/ml, in chloroform, Avanti) to a 25 ml spherical flask, yielding a molar ratio of POPC:POPG = 9:1. The spherical flask was attached to a rotary evaporator (Buchi Rotavapor, speed 8), and the lipids were dried under argon gas at room temperature for ∼30 min. The flask was then transferred to a vacuum dessicator overnight. The dried lipid film was hydrated with ∼10 ml liposome buffer using gentle swirling, and the suspension was then sonicated at a frequency of 40 Hz for 20-30 min. To generate 400 nm unilamellar liposomes, the solution was extruded 11x using a LiposoFast Basic extruder (Avestin Inc.). The extruder membrane was then changed from a pore size of 400 to 200 nm, and the solution was extruded 5x to generate 200 nm liposomes. The phospholipid concentration of the lipid suspension was quantified before reconstitution as described below. Due to losses during extrusion, the concentration was typically ∼ 3.6 mM. The liposomes were stored at 4 °C and could be used for ∼1 week after preparation.

### Quantitation of phospholipids

Phosopholipid quantitation was performed by subjecting the extruded liposomes and also proteoliposomes to oxidation by perchloric acid (Ploier and Menon, 2016 ref). Briefly, 300 µl of perchloric acid was added to a 50 µl sample, which was subjected to vortex mixing and then heated at 155 °C for 60 min. After cooling to room temperature, an aliquot (1 ml) of water was added followed by 400 µl of freshly prepared 12 g/L ammonium molybdate and 50 g/L sodium ascorbate. After vortex mixing the sample was heated at 100 °C for 10 min. The sample was then cooled to room temperature, and the absorbance at 797 nm was measured. Absorption standards of Na_2_HPO_4_ were run in parallel to generate a calibration curve.

### Reconstitution of hSERINC proteins into liposomes

Multiple reactions of 1 ml volume were typically performed simultaneously with volumes scaled accordingly. For each 1 ml reaction, reconstitution of hSERINC’s, and control proteins (A_2A_AR and Glt_Ph_) into liposomes was performed using 2 ml plastic Eppendorf tubes. An aliquot of the extruded lipid solution (typically 800 µl) was added to 34.4 µl of 10% (w/v) DDM in liposome buffer, yielding a total volume of 840 µl. The solution was incubated for 3 hrs at room temperature using an end-over-end rotator. During the last hour, 9.4 µl of the NBD-lipid label (PS, PC or PE, Avanti, Inc.) in chloroform was dried under argon and resuspended in 45 µl of liposome buffer containing 0.1% DDM. The final DDM concentration in the 1 ml reaction was ∼7 mM. For preparation of 1 ml of *empty liposomes*, 45 µl of the NBD-lipid label in 0.1% DDM liposome buffer was added to the swelled lipid solution (840 µl), to which 60 µl of 0.1% DDM liposome buffer was added, followed by 55 µl of detergent-free liposome buffer. For preparation of 1 ml of *proteoliposomes*, hSERINC3, 2 and 5 (wt, F397L, S328I) in DDM/CHS was concentrated to 1.0-1.2 mg/ml using a GE Healthcare Vivaspin (100 kDa cut-off). The protein was first diluted in 0.1% DDM liposome buffer to 150 ng/µl. The reconstitution mixture was prepared by the addition of the following solutions to 840 µl of the swelled lipid mixture in the order indicated: (1) An aliquot (x µl) of the protein, (2) 45 µl of the dissolved NBD-lipid, (3) 0.1% DDM liposome buffer (x µl) and (4) 55 µl of detergent-free liposome buffer. Liposomes and proteoliposomes were incubated for an additional hour with end-over-end rotation.

For preparation of A_2A_AR proteoliposomes, the protein was purified in a buffer containing DDM/CHS and 100 µM of the agonist adenosine (see below for details). Extruded lipids were swelled using 7.98 mM DDM for 3 hrs with end-over-end rotation. DDM/CHS solubilized A_2A_AR, NBD-PC and adenosine were added to final 1.0 ml reaction containing 7.0 mM DDM and 100 µM adenosine. Empty liposomes with adenosine were reconstituted in parallel.

For preparation of Glt_Ph_ liposomes, the purified protein (a kind gift from Olga Boudker (Weill Cornell)) was diluted in 0.1% DDM liposome buffer containing L-asp (1 mM) to 150 ng/µl,and the final 1.0 ml reaction containedincluded 1mM L-asp.

Detergent removal and generation of proteoliposomes was achieved using polystyrene beads (Bio-Beads in SM-2 Adsorbent Media, BioRad, Inc.). The protocol was adapted and refined from ^20^, so that the NBD-glucose assay (see below) demonstrated minimal leakage of the liposome- incorporated fluorescent label. To prepare ∼12 proteoliposome samples, 6-7 g of Bio-Beads were weighed into a conical tube and washed as follows: (1) Methanol (30 ml) was added to the conical tube, and the beads were incubated for 10 min with end-over-end rotation for 10 min, (2) After settling of the beads, the methanol was decanted, (3) Steps (1) and (2) were repeated 2 additional times, (4) 50 ml water was added to the beads, which were incubated as before for 10 min; (5) The water was decanted, and the beads were washed for a final 10 min in 50 ml of liposome buffer, (6) The buffer was decanted, and a glass Pasteur pipette was used to remove any residual liposome buffer and (7) The beads were stored at 4 °C.

The 1 ml proteoliposome solution was transferred to a 2.0 ml Eppendorf tube containing 100 mg of Bio-Beads, which was incubated with end-over-end rotation for 1 hr at room temperature. An additional 160 mg of Bio-Beads were then added, and the mixture was again incubated with end-over-end rotation for an additional 2 hrs. The proteoliposome solution was then transferred to a third Eppendorf tube containing 160 mg of Bio-Beads, which was incubated overnight at 4 °C. The following morning the solution was transferred to a 4^th^ Eppendorf tube containing 160 mg of Bio-Beads and incubated for a final 2 hrs at 4 °C. Any residual BioBeads were removed by passage of the solution through a Micro Bio-Spin chromatography column (0.8 ml; Bio-Rad). The proteoliposome sample in the 2 ml Eppendorf tube was placed on ice in preparation for the lipid flipping assay.

### Quantitation of protein incorporated into liposomes

Efficiency of incorporation of protein into liposomes (at the 1.5 µg/mg of lipid concentration) was determined by fluorescent imaging of Simply Blue stained proteins separated by SDS-PAGE, using purified proteins (i.e., SERINCs, A_2a_AR and Glt_Ph_) to generate a standard curve. Proteoliposomes were collected by high-speed centrifugation, solubilized with SDS sample buffer, and separated by SDS-PAGE using 4-20% gradient Mini-PROTEAN TGX gels (BioRad). In order to control for liposome related artifacts in protein quantitation, samples for standard curves were prepared by centrifugation of equivalent volumes of empty liposomes, followed by resuspension of the pellet with addition of increasing concentrations of purified protein. After a 15-minute incubation at room temperature, SDS sample buffer was added to the liposome-standard mixture prior to electrophoresis; gels were then stained with Simply Blue and destained with water. Proteins were imaged at 700 nm using an Odyssey imager (Li-Cor), and fluorescence intensity was measured. Integrated intensity of the standard curve was analyzed using GraphPad Prism, and signals from proteoliposome samples were interpolated by linear regression analysis. Typical protein recovery for hSERINC3, hSERINC5 and hSERINC2 was 43 ± 10%, 41 ± 4%, and 58 ± 11%, respectively, for proteoliposomes set up at the 1.5 µg/mg of lipid density (n=4). Phospholipid recovery for the same proteolipososomes used for protein quantitation was 81 ± 3%, 82 ± 2%, and 81 ± 1%, respectively. Errors were calculated using standard error of the mean.

### Fluorescent lipid flipping assay

Measurements were performed at 23 °C using a HORIBA Jobin Yvon Fluoromax-3 Spectrofluorometer with excitation and emission wavelengths set at 470 and 530 nm, respectively, and a slit width of 5. A 2 ml sample was prepared using 20 µl of liposomes (or proteoliposomes) and 1.98 ml of liposome buffer. The solution was pipetted into a 10 x 10 mm pathlength cuvette (Hellma High Precision Cell), inserted into the sample holder of the spectrofluorometer and stirred with a magnetic “flea” bar at speed 5. The fluorescent signal was recorded for 100 sec to ensure a stable signal prior to the addition of dithionite. Dithionite is especially prone to oxidation, and aliquots (10 mg) were weighed into 1.5 ml microfuge tubes, which were kept on ice until use. During the 100 sec equilibration scan, the dithionite solution was prepared by addition of 60 µl of ice-cold 1 M Tris (pH 10) to the microfuge tube. The dithionite was easily dissolved by vortex mixing for a few sec. At 100 sec, 40 µl of the dithionite solution was added to the 2 ml sample in the cuvette, yielding a final concentration of 20 mM dithionite.

FluoEssence v3.9 software was used to record the fluorescence vs time measurements. The plots of fluorescence intensity displayed an artifactual vertical spike at 100 sec due to light leakage while opening the lid of the sample chamber during pipetting of the dithionite solution into the cuvette. The fluorescence data were plotted on a 0 to 1 ordinate scale as normalized values, F/Fmax, where Fmax was the average fluorescence between 85-90 sec.

### NBD-glucose assay

We used an assay with water-soluble NBD-glucose^21^ to rule out the possibility that the reconstitution of hSERINC3 into liposomes resulted in permeability of the bilayer to dithionite, and thereby a greater reduction in fluorescence compared with empty liposomes (**Fig. 3b**). After the swelling step described above, an aliquot of NBD-glucose (Sigma) was added to the 1 ml reaction (instead of NBD-lipids) such that the final concentration was 60 µM NBD-glucose. In addition, to amplify any effects due to hSERINC3, the protein concentration was increased from 1.5 to 2.0 µg/mg of lipid. Although the majority of the extravesicular NBD-glucose was removed during the incubations with Bio-Beads, some residual external NBD-glucose accounted for the initial drop in fluorescence upon addition of dithionite at 100 sec. The fluorescent signal from 100-400 sec was then due to the entrapped NBD-glucose, and the nearly constant signal indicated that the addition of hSERINC3 did not elicit permeability of the dithionite through the bilayer. Lastly, the elimination of the fluorescent signal upon addition of 1% Triton X-100 at 400 sec was due to solubilization of the proteoliposomes and dithionite reduction of the entrapped NBD-glucose.

### Expression and purification of A*_2A_*AR

An amino-terminally fused T4 lysozyme–A_2A_AR construct with A_2A_AR truncated at position 316 was used as a positive control for the flippase experiments^22^. The construct contained a FLAG epitope immediately after a hemagluttinin signal sequence at the amino-terminus and a TEV cleavage site immediately following the FLAG epitope. The construct lacks the first 4 residues of the A_2A_AR amino terminus. A_2A_AR was expressed in Sf9 (*S. frugiperda*) insect cells using the Bac-to-Bac Baculovirus Expression System (Invitrogen). Sf9 cells were grown in ESF921 media (Expression Systems) at 27 °C, diluted to a density of 2.0 x 10^6^ cells/ml and infected with a high-titer baculovirus stock at a MOI of 3 for 48 hrs, harvested by centrifugation and stored at -80 °C. Viral titers were performed by the flow cytometric method^23^ on a Guava easycyte 8HT in which cells were stained with Anti-gp64-PE antibody (Expression Systems).

Membranes were prepared from receptor-infected insect cells as described^24^. All steps were performed at 4 °C unless otherwise noted. Cells were resuspended in hypotonic Buffer A (10 mM HEPES, pH 7.5, 10 mM MgCl_2_, 20 mM KCl and EDTA free protease inhibitor cocktail) and lysed by Dounce homogenization. Membranes were recovered by centrifugation at 125,000 x g and washed again with Buffer A. Two additional washes were performed with buffer A containing 1 M NaCl (Buffer B). After the final wash, membranes were weighed, resuspended in 2.5 volumes of buffer A containing 40% (v/v) glycerol (Buffer C) and flash frozen in liquid nitrogen.

For purification of A_2A_AR, frozen membranes were thawed in a 10-fold excess of 20 mM HEPES, pH 7.5, 10% glycerol, 4 mM CaCl_2_, 100 µM adenosine and EDTA free protease inhibitor cocktail (Buffer D) containing 300 mM NaCl. The suspended membranes were solubilized with 0.5% DDM/CHS for 4 hrs with gentle rotation. A_2A_AR extracts were clarified by high-speed centrifugation at 125,000 x g, and incubated with anti-FLAG M1 agarose affinity gel (Sigma-Aldrich, catalog number A4596) for overnight binding. Chromatography was then performed using a BioRad Econo-column. The bound A_2A_AR was washed with 30 column volumes of Buffer D containing 300 mM NaCl and 0.1% DDM/CHS, followed by a wash with 30 column volumes of Buffer D containing 500 mM NaCl and 0.05% DDM/CHS. The final wash was with 20 column volumes of Buffer D containing 150 mM NaCl and 0.025% DDM/CHS. The bound A_2A_AR was eluted from the FLAG-M1 affinity resin with 10 column volumes of elution buffer containing 20 mM HEPES, pH 7.5, 10% glycerol, 300 mM NaCl, 100 µM adenosine, 10 mM EDTA, 0.025% DDM/CHS and EDTA-free protease inhibitor cocktail. Eluted A_2A_AR was concentrated to 500 ml in a Vivaspin 6 mL concentrator with a 100,000 molecular weight cutoff and further purified by size exclusion chromatography using a Superdex 10 x 300 column with buffer containing 25 mM HEPES, pH 7.5, 150 NaCl, 0.025% DDM/CHS and 100 µM adenosine. Fractions containing the A_2A_AR were pooled and purified as described above, and concentrated to ∼1 mg/mL for fluorescent lipid flipping experiments.

### Cells, plasmids, and reagents

HEK293 cells were a kind gift from John A.T. Young. TZM-bl cells were obtained from the NIH AIDS Reagent Program. HEK293T cells were obtained from ATCC. Jurkat TAg and hSERINC3/5^-/-^ Jurkat TAg cells were a kind gift from Heinrich Göttlinger^25^. 293T/17 (293T) was a kind gift from Jens H. Kuhn (NIH/NIAID). HT1080-mCAT1 cells stably expressing the mCAT1, the ecotropic MLV receptor from mouse cells, have been described previously^26^. HEK293 and Jurkat cells were maintained in RPMI (Gibco) media containing 10% (v/v) fetal bovine serum, 1% Penicillin/streptomycin and 1% Glutamine. HEK293T cells were maintained in DMEM (Gibco) media containing 10% (v/v) fetal bovine serum, 1% Penicillin/streptomycin and 1% Glutamine. pNL4-3 Nef- was generated by replacing amino acids 31-33 of the Nef open reading frame with stop codons. pNL4-3 NefC contains the Nef gene from a clade C HIV-1 isolate in place of the NL4-3 Nef gene and was a kind gift from Heinrich Göttlinger. pNL4-3 *Δ*RT with V1Q3 and V4A1 tags was described previously^27^. The HIV-1 GagPol plasmid, pCMV *Δ*R8.2, was obtained from Addgene. The HIV-1 Env plasmid, pE7 NL4-3 Env, was a kind gift from Joseph Sodroski. The plasmids pCD-Env, expressing Moloney MLV Env; pBabe-Luc, an MLV vector expressing firefly luciferase; pRR1485, encoding MLV “Gag-Pol” with inactivated *env*; pRR1322, encoding MLV “Gag-Pol” with inactivated *glycogag* and *env*; pRR1321, and expressing xenotopic MLV Env were previously described^26^. A pRR1842 plasmid expressing a GFP-tagged codon-optimized MLV *gag* was constructed as follows. The codon-optimized *gag* fragment (a kind gift of Wei-Shau Hu) was cloned into pcDNA3.1(+) (Thermo Fisher) between BamHI and NotI sites. EGFP gene was amplified from pEGFP (Addgene) and then inserted into the 3‘ end of *gag* with a linker sequence "GGAGGTGGAGCATCA" in between. The CD63-mRFP plasmid was a kind gift from Gillian Griffiths^28^. hSERINC3 and hSERINC5 open reading frames were amplified from Jurkat and TZB- Bl cDNA respectively, and inserted into pcDNA3.1(+) by standard PCR techniques. Both constructs bear C-terminal FLAG tags. SERINC5 point mutants were generated by quick-change site-directed mutagenesis. The pcDNA3.1(+) hSERINC2-FLAG plasmid was a kind gift from Felipe Diaz-Griffero. GFP-Vpr was a kind gift from Tom Hope. pNL4-3 Gag-EGFP Nef- was constructed by inserting the EGFP coding sequence and a flexible poly-linker at the C-terminus of Gag.

Transmembrane protein 16F (TEM16F) is a calcium dependent phospholipid scramblase that consists of eight transmembrane *α*-helices. It is localized to the cell surface and regulates the lipid distribution across the inner and outer leaflets of the plasma membrane. It remains in an inactive state under normal physiological conditions and is only activated during cell apoptosis by the associated elevation of intracellular calcium. In its active state, TMEM16F equilibrates the phospholipid content across both leaflets of the plasma membrane by serving as an ATP-independent bidirectional lipid transporter^29^. This effectively causes elevation of the phosphatidylserine (PS) level on the outer leaflet, which is asymmetrically distributed to the cytoplasmic side in non-apoptotic cells. Here, we utilized an alternatively spliced variant form of murine TMEM16F (a kind gift from Shigekazu Nagata), which harbors an Asp-to-Gly mutation at amino acid position 409, which sensitizes TMEM16F to respond to the normal intracellular calcium concentration^30^; and a 21-amino acid-insertion at codon 24, which increases the PS scrambling activity of the D409G (now D430G) mutant. As a result, this long-form variant murine TMEM16F exhibits a constitutively high level of PS scrambling activity in an ATP- and calcium-independent manner^31^. We also observed that the WT form of mTMEM16F is not efficiently incorporated into HIV-1 particles (**Extended Data Figure 9a**). However, mTMEM16F containing the 21-amino acid-insertion at the N-terminus facilitates the incorporation into virus particles (**Extended Data Figure 9a**). Given this observation, we chose to characterize 4 mutants, which exhibit varying levels of PS exposure and inhibition of infectivity, in the context of mTMEM16F containing the N-terminal insertion. The mutants are designated as mTMEM16F DW (inactive), mTMEM16F DY (inactive), mTMEM16F GW (partially active), and mTMEM16F GY (fully active). mTMEM16F DY contains the N-terminal insertion in the context of the WT protein sequence. mTMEM16F DW contains the N-terminal insertion and the 563W mutation, the combination of which severely impairs its PS scrambling activity. mTMEM16F GW contains the 430G mutation described above, as well as the 563W mutation, which partially impairs it’s PS scrambling activity. mTMEM16F GY contains the D430G mutation described above and the wild-type 563Y residue, which confer constitutive PS scrambling activity^32^.

### Annexin V staining of retroviral particles

For HIV-1 particle staining experiments, HEK293 cells were transfected with 0.2 μg pNL4-3 Gag-EGFP *Δ*RT, 0.1 μg CD63-mRFP, 0.1 μg hSERINC/TMEM16F, and 0.05 μg GFP-Vpr plasmids in quadruplicate per 24-well culture plates using PEI. At 48 hrs post-transfection, virus particles were harvested from the supernatant, filtered through 0.45μm filters, and incubated with anti-CD63 magnetic beads (Invitrogen) for two hrs at 4°C. Virus-bead conjugates were washed with PBS +0.1% BSA, and incubated with Alexa647- conjugated annexin V diluted 1:50 in annexin V binding buffer (Invitrogen) for 45 min at ambient temperature. Samples were washed with annexin V binding buffer, and resuspended in annexin V binding buffer prior to analysis on a BD Accuri FACS instrument. Data were analyzed using FlowJo. For conditions comparing HIV-1 ±Nef, virus particles were prepared by transfecting cells with 0.8 μg pNL4-3 Nef- or pNL4-3 NefC, 0.4 μg CD63-mRFP, 0.4μg hSERINC/mTMEM16F, and 0.2 μg GFP-Vpr. Virus-bead conjugates were stained as described above and fixed with 4% PFA for 10 min at ambient temperature prior to analysis. For confocal-based experiments, HEK293 cells were plated in 6-well culture plates and transfected with 2 µg pNL4-3 *Δ*RT Nef-, 100 ng GFP-Vpr, and 100 ng hSERINC or mTMEM16F plasmids or empty vector using Fugene 6, according to the manufacturer’s instructions. After 48 hrs, virus-containing supernatants were filtered and placed in poly-lysine-coated glass-bottom dishes (MatTek). Particles were then washed and stained with Alexa594-annexin V (Thermo Fisher Scientific) in annexin binding buffer (10 mM HEPES, pH 7.4, 140 mM NaCl, 2.5 mM CaCl_2_). Confocal imaging was performed using a Nikon Eclipse TE2000-E microscope equipped with 444-, 488-, and 561-nm lasers, a Yokogawa CSU 10 spinning disc confocal laser scanning unit, and an Andor Zyla sCMOS camera. Images were quantified manually using ImageJ by drawing 20-pixel circular regions of interest around each virus particle, subtracting background fluorescence, and calculating the ratio of annexin fluorescence to GFP-Vpr fluorescence. For Jurkat-derived virus particle analysis, HEK293 cells were transfected with pNL4-3 Nef- and pNL4-3 Gag-EGFP Nef- at a ratio of 1:1. After 48 hrs, supernatants were filtered and used to spinocculate Jurkat cells at 1200 x g for 2 hrs at room temperature in the presence of polybrene. Jurkat cells were incubated for 24 hrs, washed, and cultured for 4 days. Jurkat supernatants were then collected, filtered, and subjected to microvesicle depletion using anti-human CD45 magnetic beads (BioLegend) and a separation magnet (Miltenyi Biotec). Virus particles were immobilized, stained and imaged as described above. MLV-Xeno and -Eco pseudovirions with or without GlycoGag protein were produced by transient transfection of 293T cells using Mirus TransIT-293 transfection reagent. Cells (6 x 10^5^ cells per well) in a 6-well plate were cotransfected with a mixture of Env-defective MLV “Gag-Pol” with or without GlycoGag protein (3 μg of pRR1485 or pRR1322), pBabeLuc (0.5 μg), an Env expression plasmid (0.5 μg of pCD-Env or pRR1321), pBJ5-Ser5 or pRR1839 (0.1 μg), and pUC-CMV as a filler plasmid. The GFP-tagged MLV particles were prepared in a similar way but with the addition of pRR1842 (0.15 μg) to the transfection. All supernatants were collected at 48 and 72 hrs post-transfection, pooled, and filtered through 0.22 μm filters. Filtered supernatant containing GFP-tagged MLV particles produced in the presence or absence of hSERINC5 or constitutively active mTMEM16F was placed in poly-lysine-coated glass-bottom dishes (MatTek). Particles were then washed and stained with Alexa594-annexin V (Thermo Fisher Scientific) in annexin binding buffer (10 mM HEPES, pH 7.4, 140 mM NaCl, 2.5 mM CaCl2). Confocal imaging was performed using a Leica TCS SP8 confocal laser scanning microscope (Leica, Germany). Images (n=5) of each treatment were acquired by using the same laser power and digital gain parameters with the same microscope. 12-bit images with a 1024 x 1024 field size were acquired with 60x objective lenses. After acquiring, the images were processed with ImageJ and further analyzed with CellProfiler^33^ to quantify the number of GFP-positive MLV particles and the number of those particles with which annexin V signal was colocalized. Finally, the percentage of phosphatidylserine-positive MLV particles was determined. GraphPad Prism 9 was used to perform two-way or one-way analysis of variance (ANOVA) to assess statistically significant differences.

### Single-molecule Förster resonance energy transfer microscopy (smFRET)

Labeled virus particles were prepared, imaged, and analyzed as described previously^27^. Briefly, HEK-293T cells were tranfected with 5.85 μg pNL4-3 *Δ*RT Nef- and 0.15 μg pNL4-3 *Δ*RT V1Q3/V4A1 Nef-, and 100 ng of the indicated hSERINC/mTMEM16F plasmid, using PEI (Polysciences). Viruses were collected at 48 hrs post-transfection, filtered using 0.45 μm filters, and sedimented through 15% sucrose cushions at 25,000 x g for 2 hrs at 4°C in an SW-28 swinging-bucket rotor. Virus particles were then resuspended in 1X SFP labelling buffer, containing calcium and magnesium, and incubated overnight at ambient temperature with LD550-cadaverine and LD650-CoA (Lumidyne), Transglutaminase (Sigma), and Acyl carrier protein synthetase, AcpS (homemade). Viruses were then biotinylated with DSPE-PEG2000-Biotin lipid (Avanti Polar Lipids), and purified by sedimentation through a 6-18 % continuous OptiPrep gradient at 40,000 x g for 1 hour at 4°C in an SW41 swinging-bucket rotor. Virus particles were placed on a streptavidin-coated fused silica slides and incubated in smFRET imaging buffer containing an oxygen-scavenging system composed of protocatechuic acid (PCA) and protocatechuate dioxygenase (PCD). Data were acquired on a home-built prism-TIRF microscope. Data were analyzed using a custom Matlab-based analysis software package, SPARTAN^34^, courtesy of Scott Blanchard, and custom Matlab scripts, developed in the Mothes lab. Dynamic molecule traces were combined into population FRET histograms and fitted to a 3-state Gaussian distribution centered at ∼0.15, ∼0.35 and ∼0.6 FRET.

### Virus infectivity and Western immunoblotting

For HIV-1 infectivity measurements, HEK293 cells were transfected with 0.2 μg pNL4-3 Nef- plasmid, 0.1 μg pHIV-In-GLuc plasmid, and 0.1 µg of the indicated hSERINC/mTMEM16F plasmid in quadruplicate per 24-well culture plates using PEI. Virus-containing supernatants were harvested at 48 hrs post-transfection, pooled, filtered, and used to infect TZM-Bl indicator cells. Secreted Gaussia Luciferase activity was measured 48 hrs post-infection. To measure the infectivity of the MLV pseudovirions, we first seeded the HT1080-mCAT1 cells in 12-well plates at 1 x 10^5^ cells per well. The following day, cells were pre-treated with 20 μg/ml DEAE-dextran (Sigma) at 37 °C for 30 min and then infected with MLV-Xeno or Eco pseudovirions that had been produced in the presence or absence of hSERINC5 or constitutively active mTMEM16F. At 48 h post-infection, firefly luciferase activity was measured in cell lysates, using the luciferase assay system (Promega) following the manufacturer’s protocol. Meanwhile, the amount of input virus in the filtered virus supernatant was determined by quantitative anti-p30 immunoblotting. The specific infectivity was finally determined by normalizing the luciferase signal with the amount of virus input. For Western immunoblot analysis of HIV-1 particles, cells were transfected with 2 μg pNL4-3 Nef- plasmid and 200 ng hSERINC/TMEM16F plasmids. Virus-containing supernatants were collected at 48 hrs post-transfection, filtered, and pelleted by centrifugation at 20,000 x *g* for 90 min at 4°C. Cells and virus pellets were lysing for 20 min on ice in hSERINC lysis buffer^35^ (10 mM HEPES, pH 7.5, 100 mM NaCl, 1 mM TCEP [Tris(2-carboxyethyl)phosphine], 1% DDM [n-Dodecyl-β-D-maltoside]) containing cOmplete mini protease inhibitor cocktail (Sigma-Aldrich, St. Louis, MO, USA). Lysates were mixed 1:1 with 2X LDS sample buffer containing 50 mM TCEP (Invitrogen), incubated for 5 min at ambient temperature, and subjected to SDS-PAGE. Proteins were transferred to PVDF membranes and subjected to immunoblotting with anti-HIV Ig polyclonal serum (NIH ARRRP) and anti-FLAG monoclonal antibody (Sigma-Aldrich, Clone M2). For immunoblotting of MLV particles, virions were concentrated by ultracentrifugaton (25,000 x *g*, at 4 °C for 1.5 hrs) through a cushion of 20 % sucrose prepared in PBS (Invitrogen). The pellets were resuspended in PBS, and the virus samples for Western blot analysis were prepared in 1 x NuPAGE LDS sample buffer containing 50 mM TCEP-HCl. To prevent hSERINC5 precipitation, samples were not heated above 37 °C. Filtered supernatant and concentrated virions underwent NuPAGE electrophoresis using 4 to 12 % Bis-Tris polyacrylamide gels (Invitrogen), followed by transfer to Immobilon-FL polyvinylidene difluoride membrane (Millipore). Membranes were blocked in Odyssey blocking buffer (Li-Cor) and probed with the following primary antibodies: rabbit anti- p30CA (NIH AIDS Reagent Program), rabbit anti-hSERINC5 (Abcam), rabbit anti-MLV gp70 (NIH AIDS Reagent Program), and rabbit anti-FLAG (Invitrogen). Secondary antibodies conjugated to DyLight 800 or 680 (Li-Cor) and the Li-Cor Odyssey imaging system were applied to specifically detect the corresponding protein. Images were analyzed using ImageStudioLite (Li-Cor).

### Cell viability measurements

To determine the viability of the virus producing cells, 30,000 HEK293E cells were co-transfected on a 96 well plate with 14 ng of pcDNA3.1-based DNA constructs expressing either wild type or mutant hSERINC5 or mTMEM16F, along with 28 ng of pNL4-3 Gag-EGFP Nef- and 14 ng of CD63-mRFP. Forty-eight hrs post transfection, 100 µL of freshly thawed CellTiter-Glo One Solution Assay (Promega) reagents were added to each transfected 96 well and mixed well to induce cell lysis. The mixed content was then incubated at room temperature for 10 min, during which the released ATP from lysed cells activate the Luciferin to generate luminescent signals that reflect the total ATP level within and hence the viability of the transfected cells. Luminescence was then collected for each well on a TECAN SPARK^®^ Multimode Microplate Reader.

### Virus Capture Assay (VCA)

HEK-293T cells were transfected with 3.5 μg pNL4-3 *Δ*Vpr *Δ*Env F-Luc, 1 μg VSV-G, 3.5 μg pNL4-3 Nef-, 1 μg GFP, and 200 ng of the indicated hSERINC plasmid in 10 cm tissue culture dishes. Viruses were then subjected to the VCA with the indicated antibody at a concentration of 5 μg/ml, as described previously^36^.

### Molecular dynamics simulations

All MD simulation systems were prepared with the CHARMM-GUI membrane builder^37^. Three independent MD systems were prepared for hSERINC3 to sample different initial lipid positions. The lipid composition of the bilayers was selected to approximate that of an HIV infected cell outer membrane in the region of viral fusion^38^. Each bilayer consisted of approximately 230 lipid and cholesterol molecules in each leaflet. In each system, the approximate composition of the extracellular leaflet was 26% cholesterol, 17% POPC, 14% POPE, 29% POPS, and 14% sphingomyelin and the cytoplasmic leaflet was 26% cholesterol, 19%POPC, 15%POPE, 32% POPS and 8%PIP2. The systems were solvated with TIP3 water molecules and neutralized and ionized with 140 mM NaCl. Five calcium ions were added to each system to mimic physiologic calcium concentration. Each hSERINC MD system consisted of ∼155,000 atoms. The simulations were performed at constant pressure (1 atm) and constant temperature (310 K). All hSERINC3 MD simulations were run with GROMACS version 2019.4 with CUDA support on Linux workstations equipped with consumer-grade NVIDIA graphics cards.

## Extended Data Figures

**Extended Data Figure 1.**
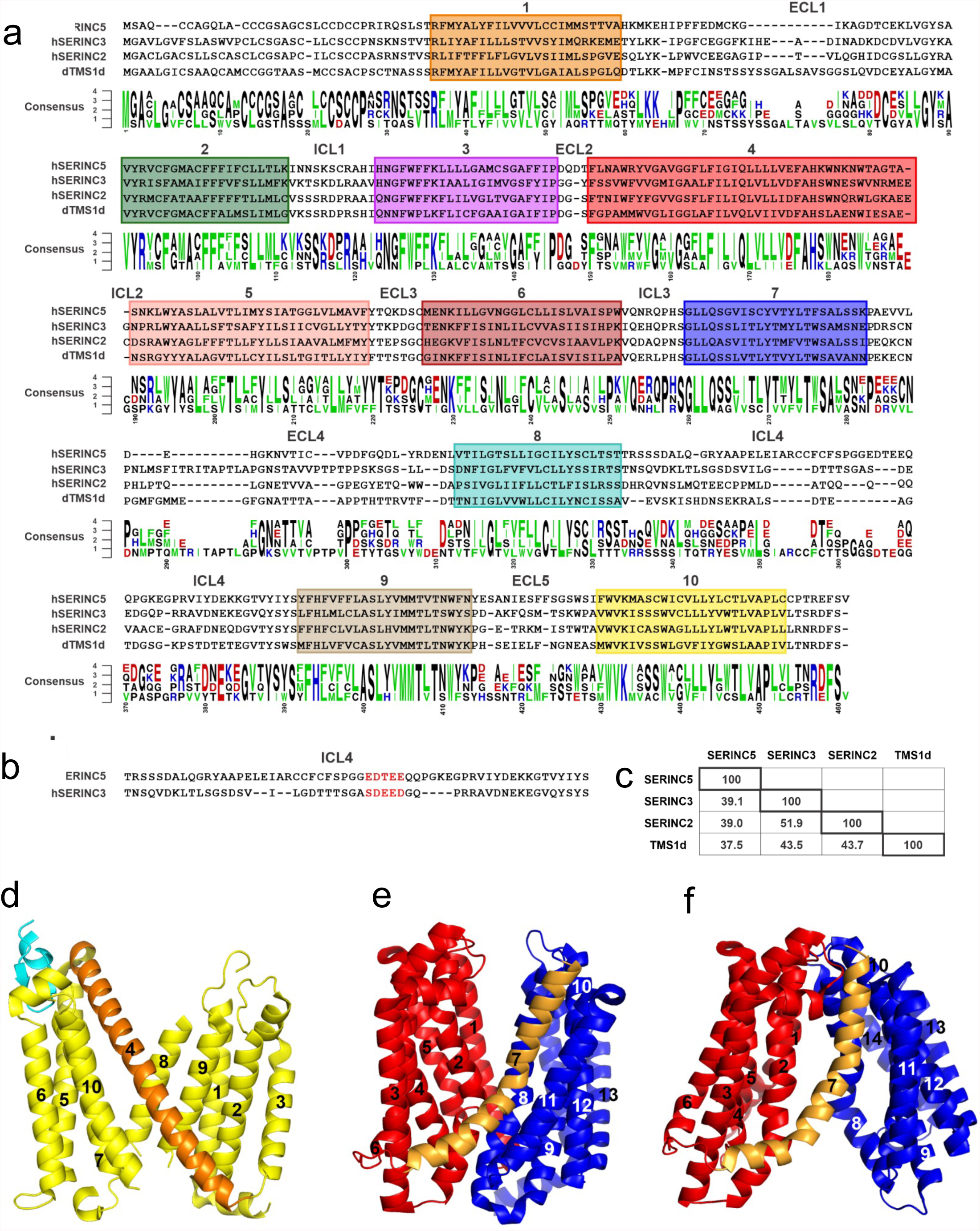
Sequence alignments and comparisons using Clustalω and architectural homology between Serinc3 and the bacterial lipid II flipase MurJ. (**a**) The sequence alignments include hSERINCs 5 and 3 and a Drosophila ortholog (TMS1d), which display restriction activity, and hSERINC2, which does not display restriction activity. All four proteins display ten predicted transmembrane *α*-helices (H1-H10). The predicted intracellular (ICL) and extracellular (ECL) loops are indicated. (**b**) Alignment of the ICL4 regions from hSERINCs 5 and 3 from (**a**), highlighting the analogous acidic cluster motifs in red. (**c**) The Percent Identity matrix of hSERINCS5, 3 and 2 and TMS1d indicate 40-50% identity between the proteins.(**d**) Structure of Serinc3 (PDB: 7RU6) colored yellow with crossmember (H4, orange) connecting the lobe comprised of helices 1, 2, 3, 8 and 9 with the lobe comprised of helices 5, 6, 7 and 10. Fragment of Fab bound is shown in cyan. (**e**) Outward open conformation of MurJ (PDB: 6NC9) with analogous crossmember (helix 7, gold) connecting lobe comprised of helices 1-6 (red) with the lobe comprised of helices 8-14 (blue). Note that notwithstanding the homologous lobe architecture between Serinc3 and MurJ, the helical composition of the lobes is not conserved. (**f**) Inward open conformation of MurJ (PDB: 6NC7), colored as in *(b*).

**Extended Data Figure 2.**
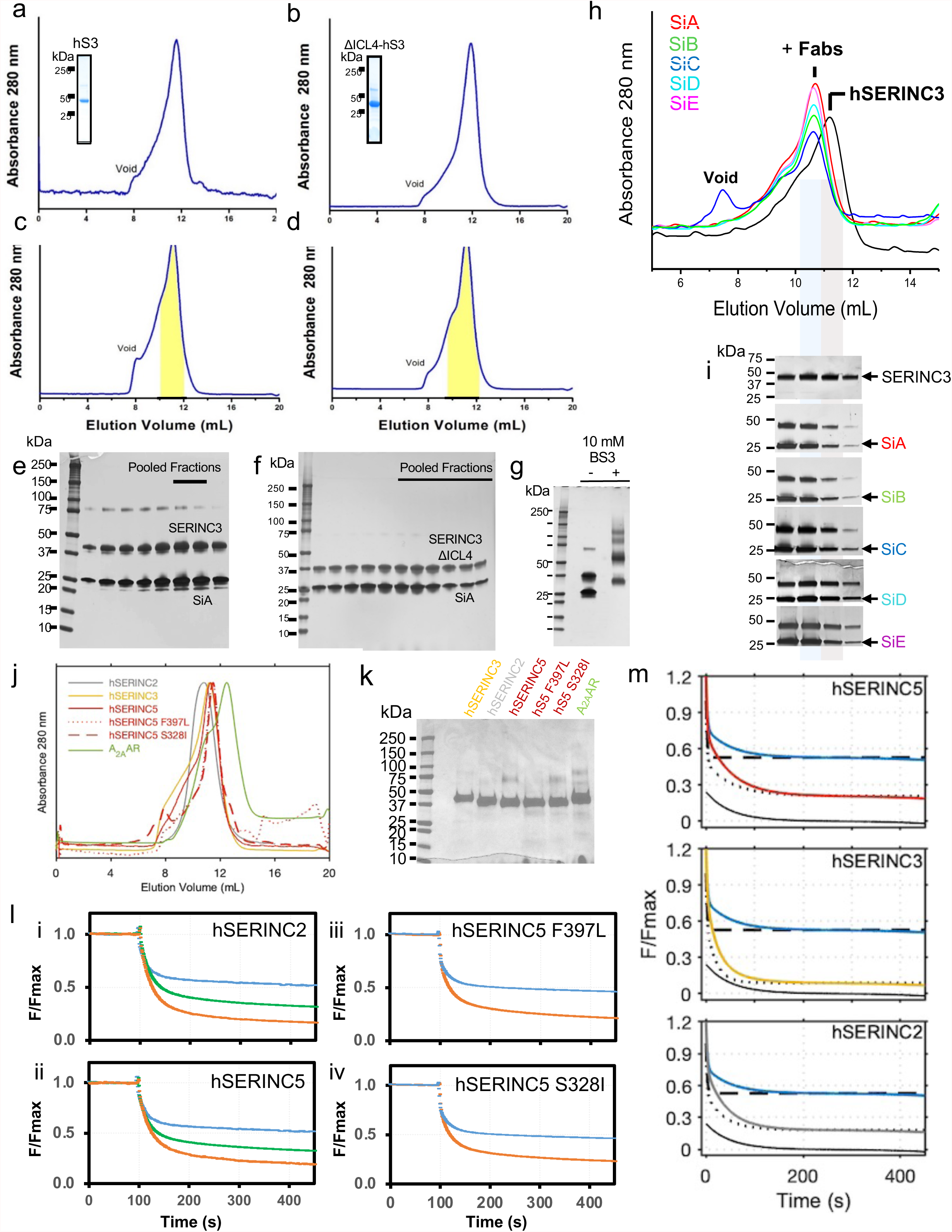
Purification of hSERINCs for structural and functional characterization. <u>Purification of WT hSERINC3, ΔICL4-hSERINC3 and formation of complexes with the Fab SiA (Kossiakoff designation)</u>. (**a-i**) Shown are the experiments used to generate purified full-length WT hSERINC3 and ΔICL4-hSERINC3 used for the single-particle cryoEM data collections that resulted in the maps shown in Figures 1 and 2. SEC chromatograms from Superdex200 gel filtration purified hSERINC3 (**a**) or ΔICL4-hSERINC3 (**b**) solubilized in DDM/CHS and exchanged into GDN. Insets show the Simply Blue gels of the purified proteins after elution from the Strep-Tactin beads that were loaded onto the Superdex200 column. Fractions with monomeric hSERINC3 (**c**) or ΔICL4-hSERINC3 (**d**) containing monodisperse protein were then incubated with the Fab SiA and subjected to further gel filtration on the Superdex200 column. The corresponding SEC peaks are shown. Fractions (highlighted in yellow) that were analyzed by SDS-PAGE and silver-stained are shown for hSERINC3-Fab SiA (**e**) and ΔICL4-hSERINC3 (**f**). hSERINC3 and ΔICL4-hSERINC3 samples were pooled as indicated (**e,f**), concentrated and used for structure determination. (**g**) The ΔICL4-hSERINC3 sample was cross-linked with 10 mM bis(sulfosuccinimidyl)suberate (BS3) prior to preparation of frozen-hydrated specimens. (**h**) SEC chromatograms displayed a higher molecular weight shift of hSERINC3 in the presence of Fabs designated SiA (red), SiB (green), SiC (blue), SiD (cyan) and SiE (magenta) in 0.02% DDM/CHS in comparison with hSERINC3 alone (black). (**i**) Silver-stained SDS-PAGE fractions across the peaks in (**h**) for hSERINC3 alone and each Fab. Peak fractions for hSERINC3 (light gray) and hSERINC3-Fab (light blue) are boxed in (**h**) and (**i**). Shown to the left of the hSERINC3 gel at the top are molecular weight standards. <u>Purification of hSERINCs and A_2A_AR for *in vitro* lipid flipping experiments</u>. (**j**) Representative SEC chromatograms from Superdex 200 gel filtration of purified hSERINC2, hSERINC3, hSERINC5, hSERINC5-F397L, hSERINC5-S328I and A_2A_AR in DDM/CHS. For hSERINC2, the chromatogram is from a second gel filtration step used to further purify hSERINC2 monomers. (**k**) SDS-PAGE of the purified proteins stained by Simply Blue after SEC gel filtration. <u>hSERINC5, hSERINC2 and hSERINC5 mutants F397L and S328I exhibit lipid flippase activity for PC</u>. (**l**) Representative fluorescence traces corresponding to dithionite treatment of NBD-PC for empty liposomes (blue) or proteoliposomes containing (i) hSERINC2 and (ii) hSERINC5, and the mutants (iii) hSERINC5 F397L and and (iv) hSERINC5 S328I. The orange curves are at a protein concentration of 1.5 µg/mg of lipid, and the green curves in i. and ii. are at 0.5 µg/mg of lipid (green). (**m**) Representative traces of exponential fits to fluorescent decay data for hSERINCs 5, 3 and 2, such as those shown in (**l**) panels (i) and (ii) and Figure 3c. The exponential “curve stripping” method was as follows. The first few seconds of the protein-free liposome data (blue) representing the quenching of fluorescence in the outer bilayer leaflet by dithionite can be fitted using a single exponential decay curve (black dashed). Subtracting the black dashed curve from the blue curve leaves a double exponential curve (black solid). This double exponential is subtracted from the observed proteoliposome decay curve, leaving the decay resulting from diothionite quenching in combination with SERINC lipid flipping (black dotted). Fluorescence decay of empty liposomes (blue) results from the sum of quenching due to dithionite reduction of lipids in the outer membrane leaflet of the liposomes (black dashed) and a slower reaction of unknown origin (black solid). Decay of proteoliposomes containing hSERINC5 (i, red), hSERINC3 (ii, gold), and hSERINC2 (iii, gray) results from the sum of the unknown slow reaction (black solid) and quenching due to dithionite reduction of fluorescence in the outer leaflet as each hSERINC flips the NBD-PC lipids (black dotted). The forward (*α*) and backward (*β*) flipping rate within each isoform is similar, supporting the assumption that there is no preferential direction for incorporation of SERINCs into liposomes. However, the flipping rates between isoforms are different. hSERINC3 flips fastest (*α*= (5.84 ± 0.15) × 10^-2^ s^-1^, *β* = (5.5 ± 0.3) × 10^-2^ s^-1^), hSERINC5 the slowest (*α* = (1.92 ± 0.06) × 10^-2^ s^-1^, *β*= (1.38 ± 0.08) × 10^-2^ s^-1^), and hSERINC2 at an intermediate rate (*α*= (2.87 ± 0.08) × 10!" s-1, *β* = (2.45 ± 0.13) × 10^-2^ s^-1^). On average for all 3 SERINC proteins, ∼77% of the fluorescence decay in proteoliposomes can be attributed to diothionite quenching of outer leaflet lipids in combination with SERINC lipid flipping, while 23% is due to other phenomena, such as leaky liposomes and/or a fraction of multilammelar liposomes.

**Extended Data Figure 3.**
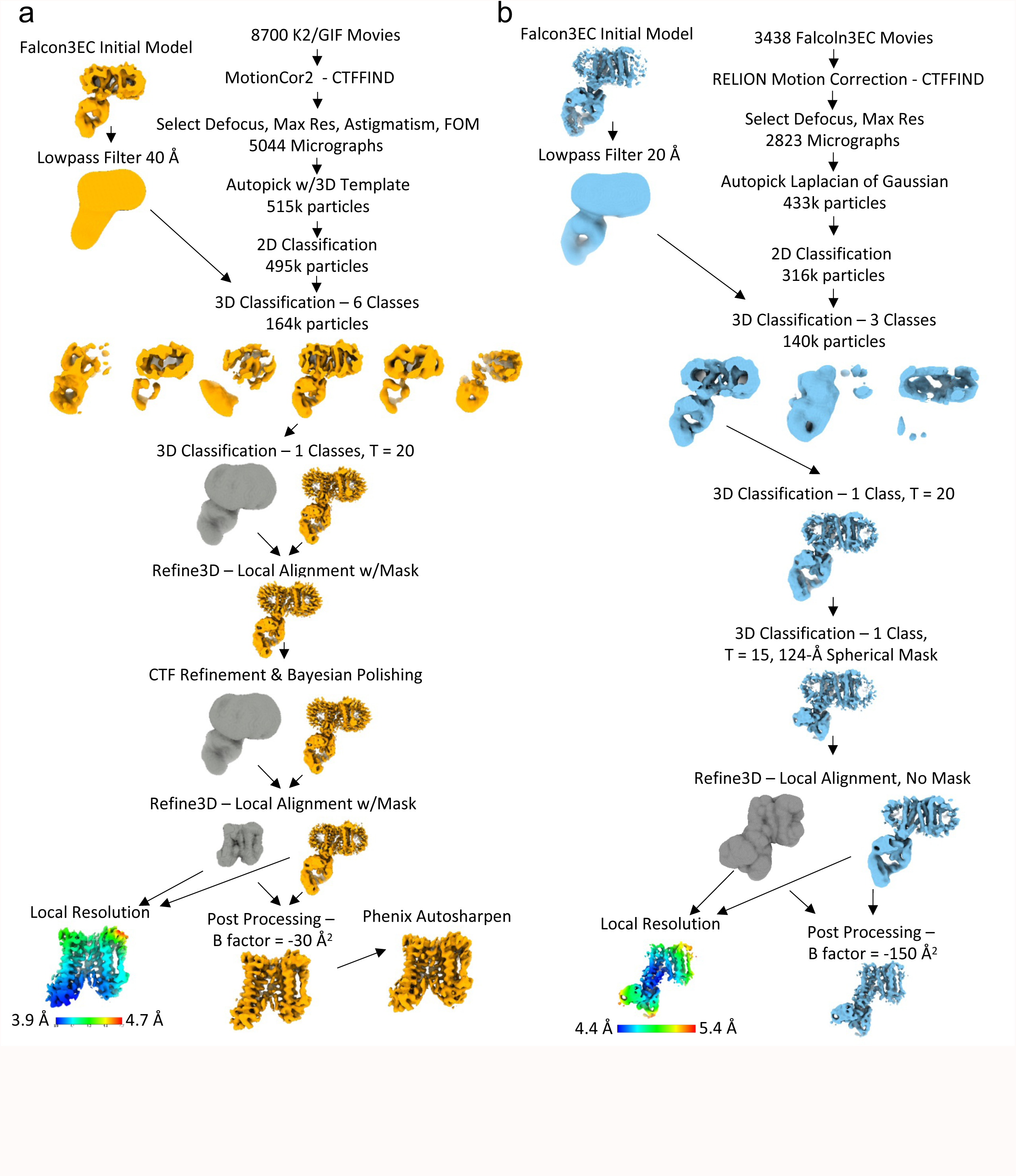
RELION processing flowcharts of the WT hSERINC3-Fab complex (a, gold) and the ΔICL4-hSERINC3-Fab complex (b, blue). Particles selected from 2D classifications were combined and further processed in 3D. An initial model for 3D had been generated from a dataset collected at the MEMC using the Falcon 3EC direct detector. The final particle set was refined using 3D auto-refine with a mask, followed by CTF refinement and Bayesian polishing, which was then followed by an additional round of 3D auto-refine with a mask. Details of the data processing strategy are described in the Methods section (dataset from NCCAT 20190102). hSERINC3 ΔICL4-Fab particles selected from 2D classifications were combined and further processed in 3D. An initial model for 3D had been generated from the hSERINC3 ΔICL4-Fab without supplements dataset that had been low-passed filtered to 20 Å. One class from the initial 3D classification was subjected to an additional round of 3D classification (T=20) followed by another round with a 124 Å spherical mask and T=15. 3D refinement with local alignment and without a mask was followed by post-processing with a B-factor of -150 Å^2^ to generate the final map. Color bars indicate the resolution of the final maps. (Dataset for this figure from MEMC 20190621.)

**Extended Data Figure 4.**
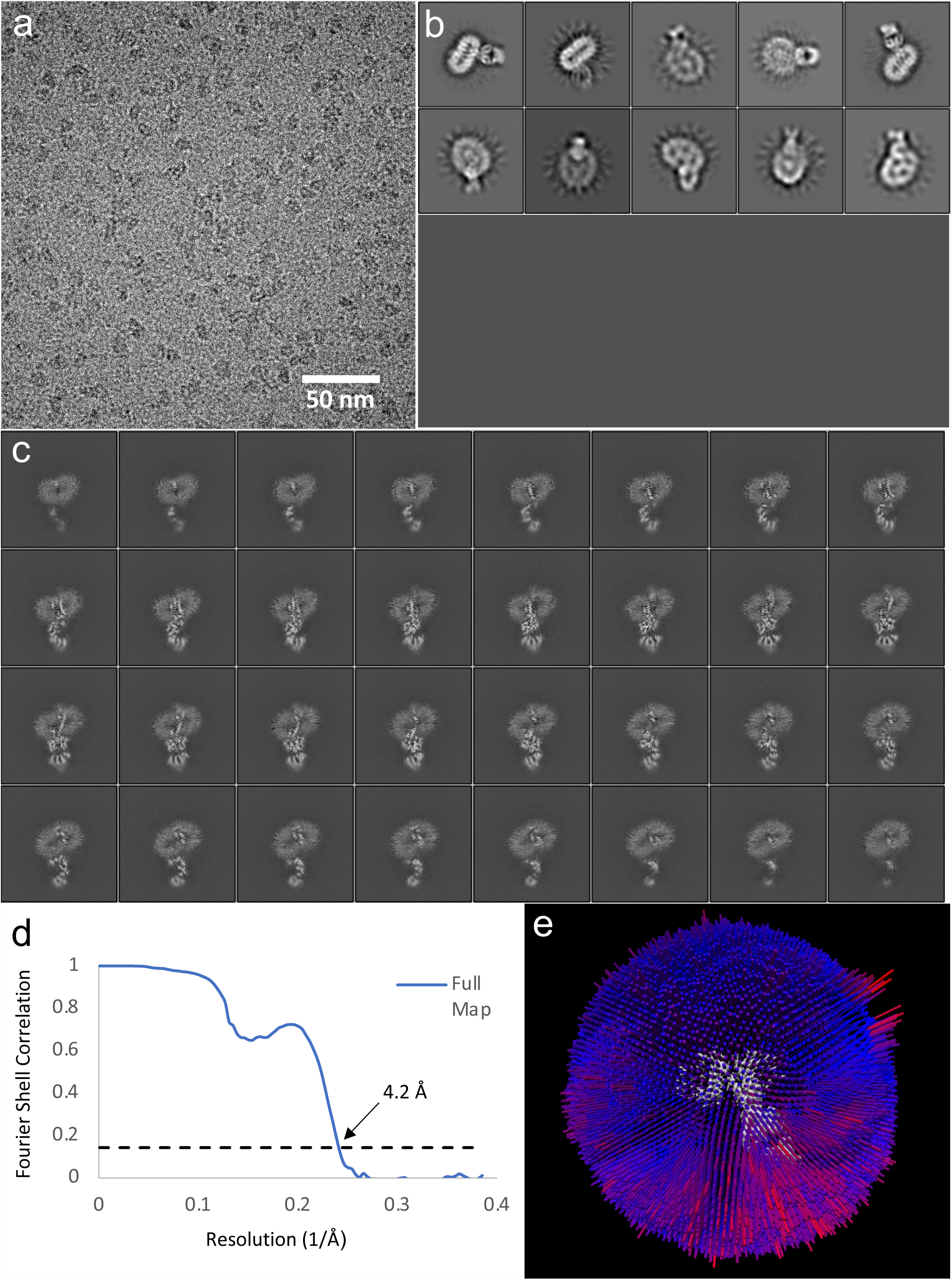
RELION processing of the wildtype hSERINC3-Fab complex. (**a**) Representative cryoEM image of hSERINC3-Fab complexes displays homogeneous and monodisperse particles. (**b**) 2D class averages of particles from cryoEM images. (**c**) 2D slices through the 3D map. (**d**) FSC (Fourier shell correlation) curve between independently refined half-maps of the final reconstruction of the hSERINC3–Fab complex. Resolution of the final map was evaluated by gsFSC (gold-standard FSC) method^39^ where the FSC curve crossed a correlation value of 0.143. (**e**) Euler angle particle distribution of reconstruction. (Dataset for this figure from NCCAT 20190102.)

**Extended Data Figure 5.**
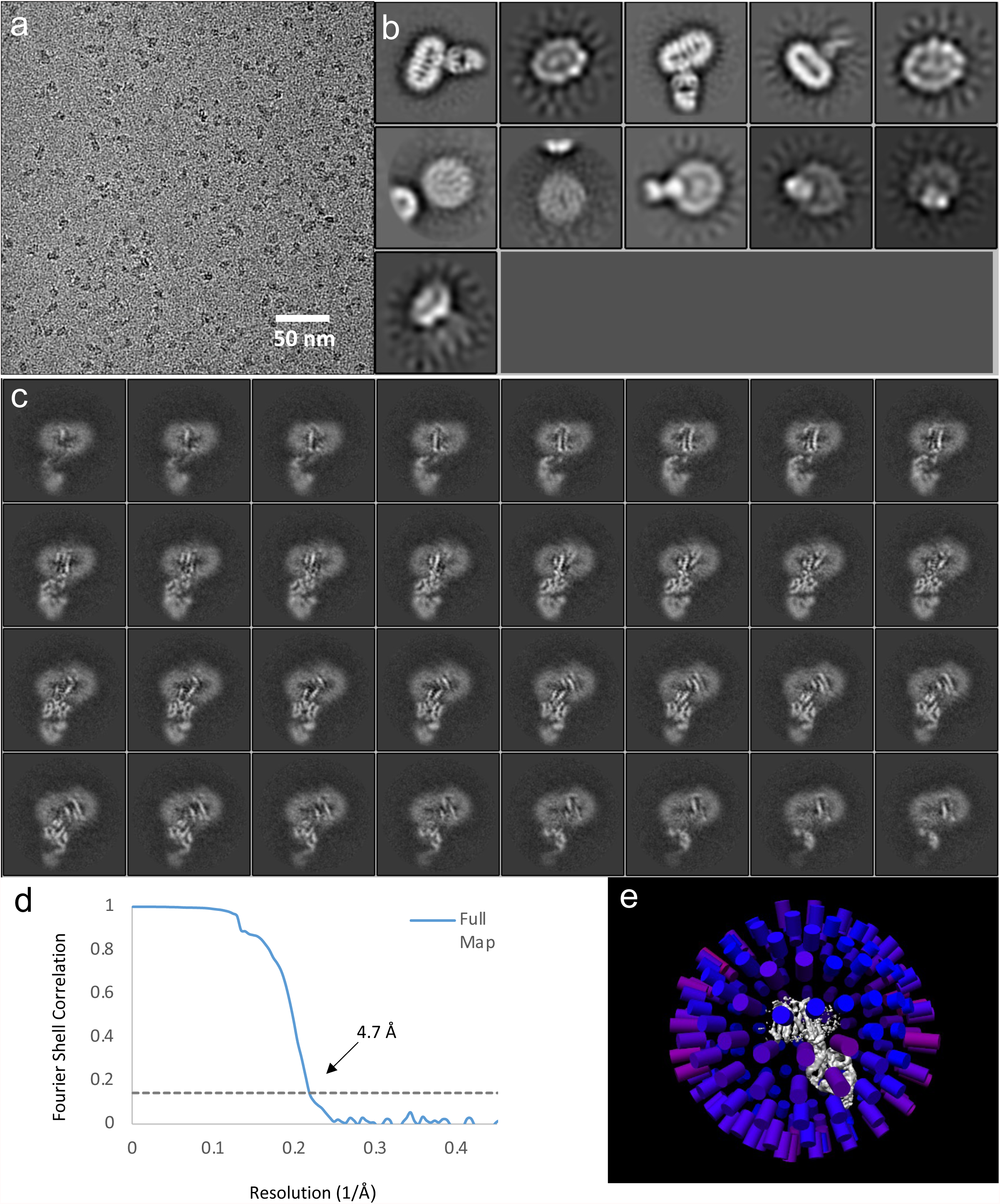
RELION processing of the ΔICL4-hSERINC3-Fab complex. (**a**) Representative cryoEM image of ΔICL4-hSERINC3-Fab complex displays homogeneous and monodisperse particles. (**b**) 2D class averages of particles from cryoEM images. (**c**) 2D slices through the 3D map. (**d**) FSC (Fourier shell correlation) curve between independently refined half-maps of the final reconstruction of the ΔICL4-hSERINC3-Fab complex. Resolution of the final map was evaluated by gsFSC (gold-standard FSC) method^39^ where the FSC curve crossed a correlation value of 0.143. (**e**) Euler angle particle distribution of reconstruction. (Dataset for this figure from MEMC 20190311.)

**Extended Data Figure 6.**
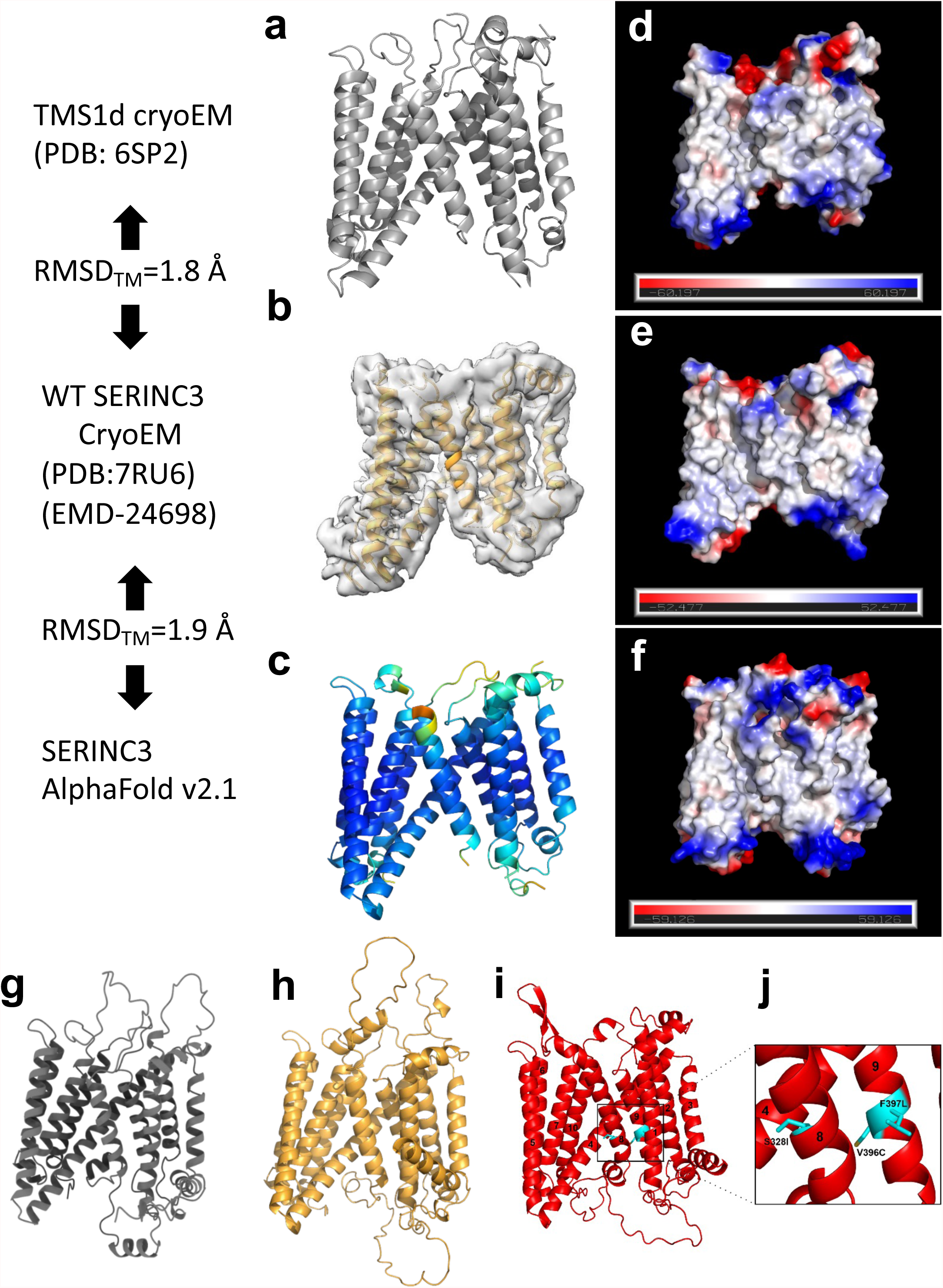
Comparison of SERINC models. **(a)** The Drosophila TMS1d monomer extracted from the cryoEM structure of the hexameric protein. (**b)** The WT hSERINC3 cryoEM model and map. (**c)** An AlphaFold model of hSERINC3 colored by pLDDT confidence score (blue, very high; cyan, high; yellow, low; orange very low). Low and very low confidence loops were the same regions missing in the cryoEM maps and were removed for comparison. Backbone RMSDs for the TM domains were calculated in PyMOL. RMSDs are lower when aligning the TM bundles separately. (**d – f)** Vacuum electrostatic surface potentials for each model calculated in PyMOL demonstrating fairly similar electrostatic distributions for the three models (red, anionic and blue, cationic). (**g**) Ribbon representation of AlphaFold 3D models for hSERINC proteins, hSERINC3 (gold), hSERINC5 (red) hSERINC2 (grey). (**g**) Ribbon representation of AlphaFold 3D model for (**g**) hSERINC2 (grey), (**h**) hSERINC3 (gold) and (**i**) hSERINC5 (red); point mutations in H8 and H9 are colored in cyan and boxed. (**j**) Closeup of (**i**) highlighting the S328I mutation in H 8 and the V396C and F397L mutations in H9; H1 was removed for clarity.

**Extended Data Figure 7.**
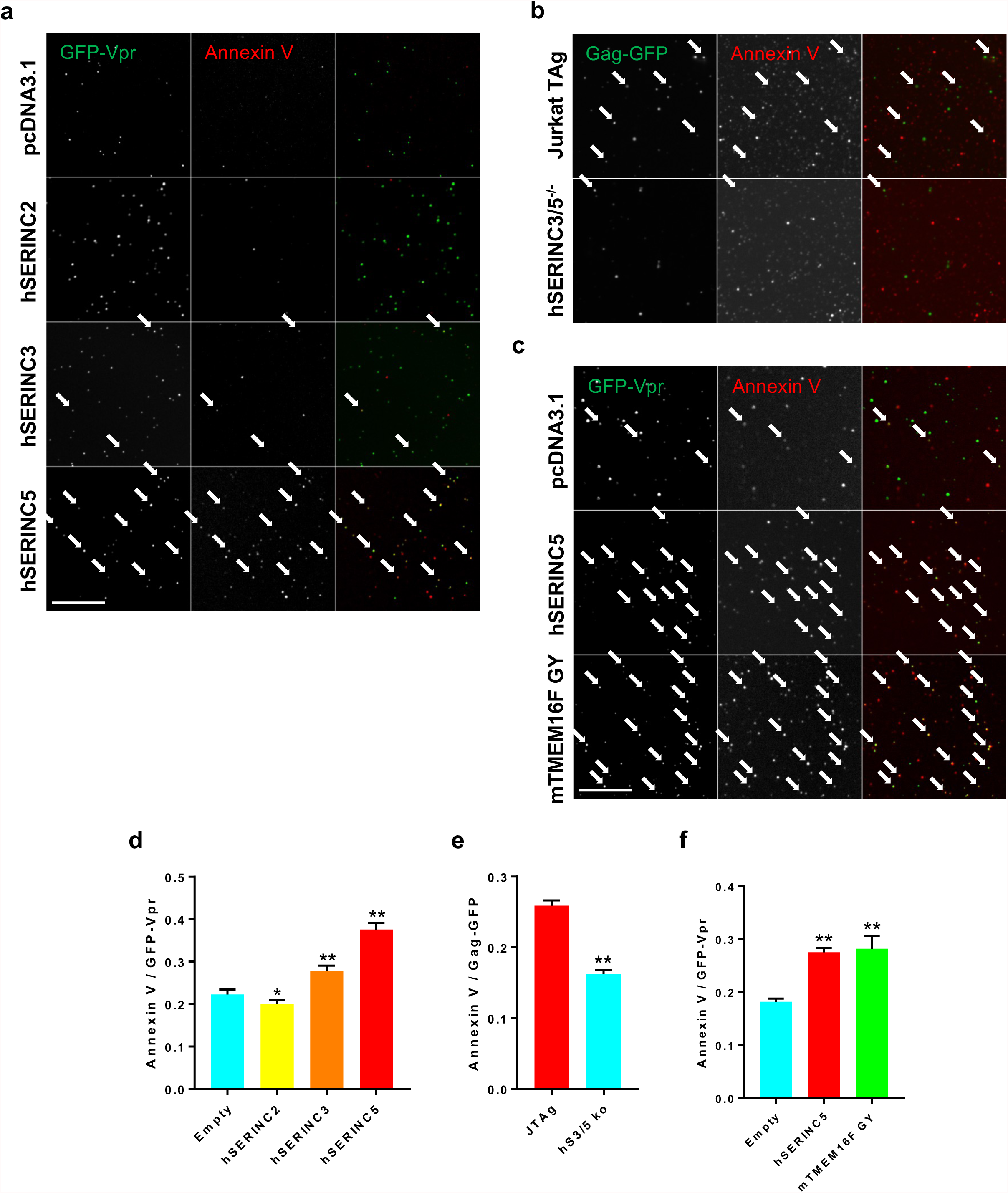
hSERINC3, hSERINC5, and mTMEM16F GY expose PS on the surface of HIV-1 particles. (**a**) HEK293-derived HIV-1_NL4-3_*Δ*RT*Δ*Nef virus particles containing GFP-Vpr were produced in the presence of the indicated hSERINC plasmid, immobilized on poly-lysine coated coverslips and stained for PS exposure using Alexa594 annexin V. (**b**) HIV-1_NL4- 3_*Δ*Nef virus particles containing Gag-EGFP were produced from parental or hSERINC3/5 knockout Jurkat TAg cells, immobilized on poly-lysine coated coverslips, and stained with Alexa594 annexin V, prior to fixation with 4% PFA, and analysis by confocal microscopy as in (**a**). (**c**) HIV-1_NL4-3_*Δ*RT*Δ*Nef virus particles containing GFP-Vpr were produced in the presence of WT hSERINC5 or mTMEM16F GY plasmid, stained, and analyzed as in (**a**). (**d-f**) Mean annexin V fluorescence for individual HIV-1 particles was quantified and normalized to GFP intensity. Particles (n=400) were quantified per condition, from two independent experiments. Scalebar = 5mm. Values represent mean annexin V intensity per GFP-Vpr or Gag-GFP intensity ± SEM. A non-parametric Mann-Whitney test was used to determine significance; * p < 0.05, ** p < 0.01.

**Extended Data Figure 8.**
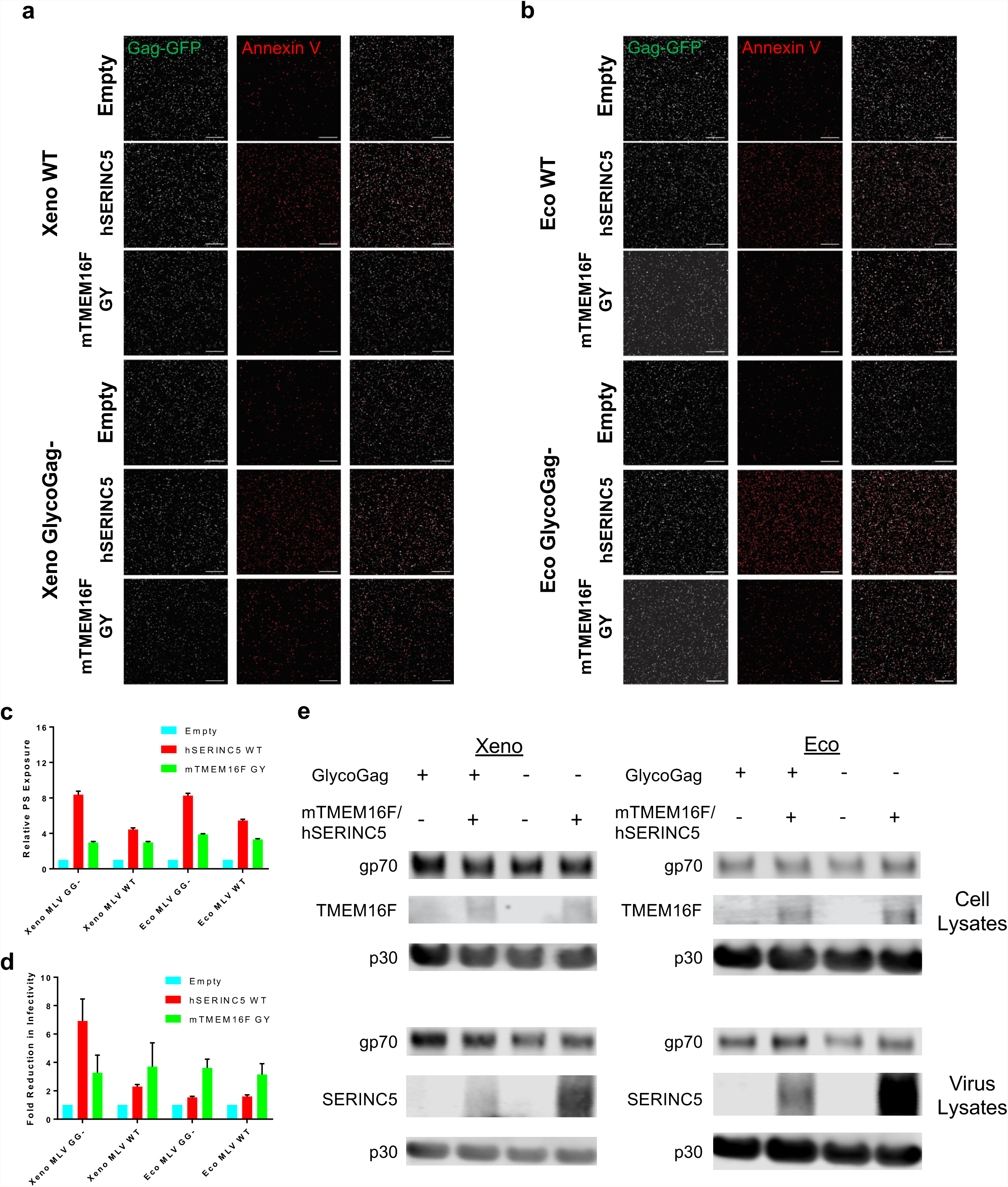
hSERINC5 and mTMEM16F GY expose PS on the surface of MLV particles. (**a**) MLV VLPs containing GagGFP and Xenotropic MLV Env with or without GlycoGag were produced in the presence of the indicated plasmid. VLPs were immobilized on poly-lysine coated coverslips, stained with Alexa594-annexin V, and imaged by confocal microscopy. (**b**) MLV VLPs containing Eco MLV Env were produced and analyzed as in (**a**). Scalebar = 10 µm. (**c**) Percent annexin V-positive VLPs were quantified using Cellprofiler. Values represent the mean of 5 independent experiments ± SEM. (**d**) Relative infectivity of viruses used in (**a**) and (**b**) was determined by firefly luciferase assay in HT1080-mCAT1 cells. Values shown represent the mean of 3 independent experiments ± SEM. Scalebar = 10 μm. (**e**) MLV VLPs and virus-producing 293T cells were produced as in (**a**) and (**b**), lysed, and subjected to SDS-PAGE and Western immunoblotting for MLV Env (gp70), MLV CA (p30), and hSERINC5/mTMEM16F GY (using anti-FLAG antibody). Xeno – xenotropic MLV Env, Eco – ecotropic MLV Env.

**Extended Data Figure 9.**
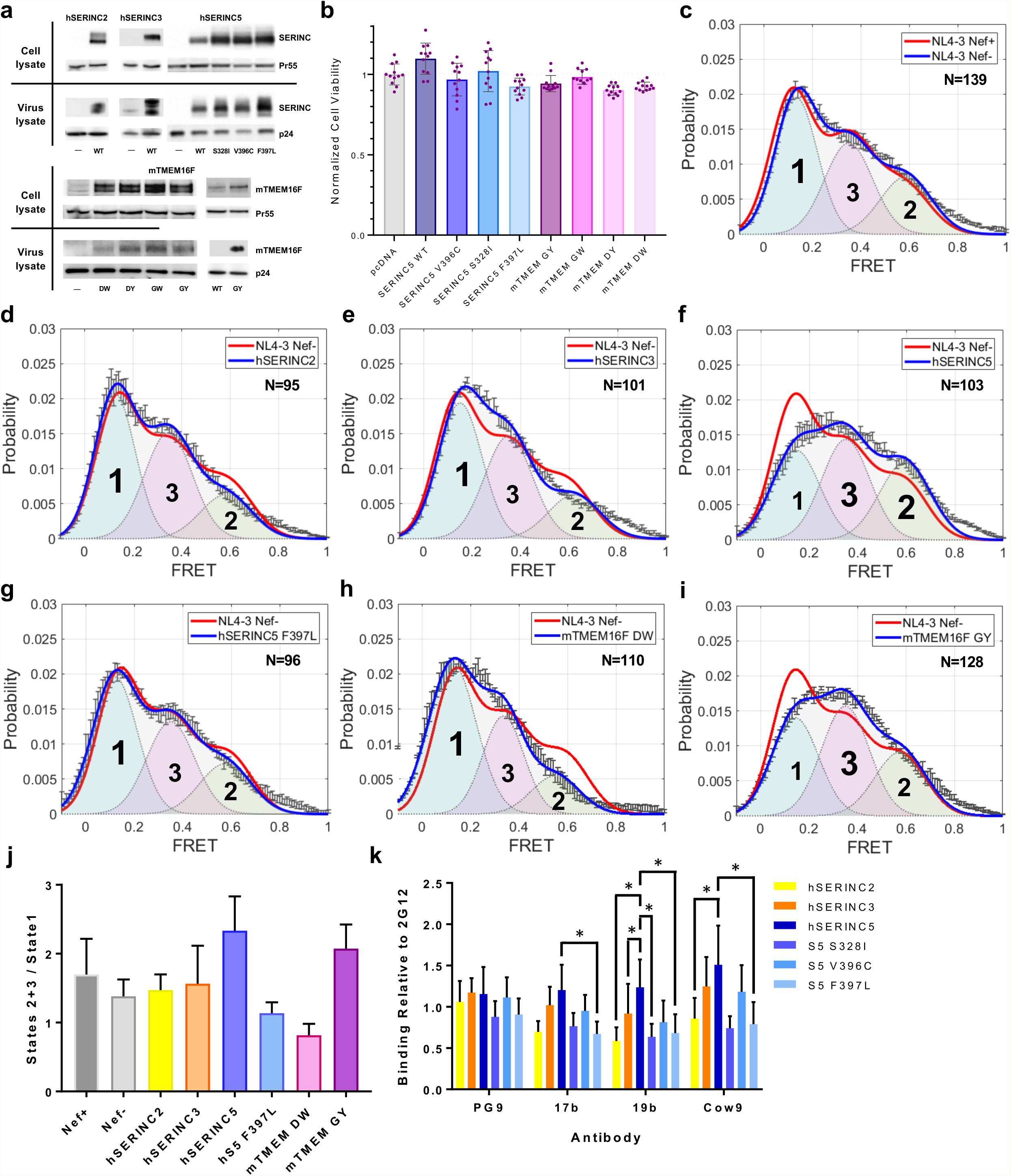
hSERINC5 and mTMEM16F GY expression promotes opening of the HIV-1 Env trimer. (a) HIV-1_NL4-3_ΔNef viruses were produced in the presence of the indicated hSERINC or mTMEM16F plasmid. Cell and virus lysates were subjected to SDS-PAGE and Western blotting for HIV-1 Gag (Pr55), HIV-1 CA (p24), and the indicated hSERINC or mTMEM16F protein (using anti-FLAG antibody, except for hSERINC3, which was detected with rabbit anti-SERINC3 antibody). (b) HEK293 cells were transfected with HIV-1_NL4-3_ΔRTΔNef and the indicated hSERINC5/mTMEM16F plasmid. Cell viability was assessed by measuring the total ATP content of freshly-prepared cell lysates using a commercially available assay kit. (c-i) HIV- 1_NL4-3_ΔRTΔNef virus particles, containing V1Q3 and V4A1 tags in gp120, were produced in the presence of the indicated hSERINC/mTMEM16F plasmids. Virus particles were enzymatically labeled with Cy3 (donor) and Cy5 (acceptor) fluorophores, purified, and analyzed by single-molecule Förster-resonance energy transfer (smFRET). A comparison between Nef+ and Nef-virus particles is shown in (c). All other experimental results are presented in relation to Nef-. Conformational states are designated as State 1 (pre-triggered, closed conformation), State 2 (necessary, partially-open, intermediate conformation), and State 3 (fully-open, CD4-bound conformation) The total number of raw FRET traces used to construct each histogram is shown as N). p values were calculated from the sum of State 2 and State 3 occupancies divided by State 1 occupancy from 3 technical replicates. (j) Occupancies for each conformational state were calculated as the area under each Gaussian curve. The ratio of open to closed Env trimers is displayed as the ratio of State 2 occupancy + State 3 occupancy / State 1 occupancy. (k) HIV-1_NL4- 3_ VLPs were prepared and subjected to virus capture assay (VCA) using the indicated antibody. All results are displayed as the amount of virus binding relative to 2G12. Values represent the mean of 5 independent experiments ± SEM. Asterisks identify statistically significant (p < 0.05) differences in the comparisons: 17b Antibody: WT hSERINC5 shows greater antibody binding than the hSERINC5 mutants S328I, V396C and F397L; 19b Antibody: WT hSERINC5 shows greater antibody binding than WT hSERINC3, WT hSERINC2 and hSERINC5 mutants S328I and F397L (but not V396C); Cow9 Antibody: WT hSERINC5 shows greater antibody binding than WT hSERINC2 (but not hSERINC3) and hSERINC5 mutant F397L (but not S328I and V396C)

**Extended Data Table 1.** CryoEM data collection, refinement and validation statistics. Summary of relevant parameters used during cryoEM data collection and processing. Refinement and validation statistics are provided for the molecular model of WT hSERINC3-Fab and ΔICL4- hSERINC3-Fab crosslinked with BS3. *Data collected at NCCAT; # data collected at the UVa Molecular Electron Microscopy Core.

| | WT hSERINC3-<sup>*</sup><br>Fab | ΔICL4-hSERINC3-<sup>#</sup><br>Fab + BS3 |
| --- | --- | --- |
| **Data collection and processing** |  |  |
| Voltage (kV) | 300 | 300 |
| Energy filter (eV) | 20 |  |
| Electron exposure (e<sup>-</sup>/Å<sup>2</sup>) | 44 | 58 |
| Defocus range (- μm) | 0.8 - 1.5 | 1.0 - 2.2 |
| Pixel size (Å) | 0.56 | 0.67 |
| Total micrograph movies | 8710 | 1574 |
| Selected micrograph movies | 5044 | 1418 |
| Initial particle images (no.) | 495,000 | 247,000 |
| Final particle images (no.) | 164,000 | 121,000 |
| Map resolution (Å) (Fab-proximal/Fab-distal) | 4.2 (3.6/4.4) | 5.1 |
| Map sharpening B factor (Å<sup>2</sup>) | -30 | -150 |
| FSC threshold | 0.143 | 0.143 |
| **Refinement** |  |  |
| Refinement resolution (Å) | 3.7 | 4.0 |
| Model composition |  |  |
| Protein residues | 303 | 302 |
| Nonhydrogen atoms | 4859 | 4940 |
| B factors (Å<sup>2</sup>) (min - max) | 147 (106 - 187) | 262 (184 - 364) |
| **Protein** |  |  |
| r.m.s. deviations |  |  |
| Bond lengths (Å) | 0.004 | 0.011 |
| Bond angles (°) | 0.8 | 1.3 |
| **Validation** |  |  |
| MolProbity score | 1.53 | 1.55 |
| Clashscore | 2.68 | 3.85 |
| Poor rotamers (%) | 0 | 0.37 |
| **Ramachandran** |  |  |
| Favored (%) | 92.2 | 94.5 |
| Allowed (%) | 9.7 | 5.5 |
| Disallowed (%) | 0 | 0 |

**Extended Data Movie 1.**
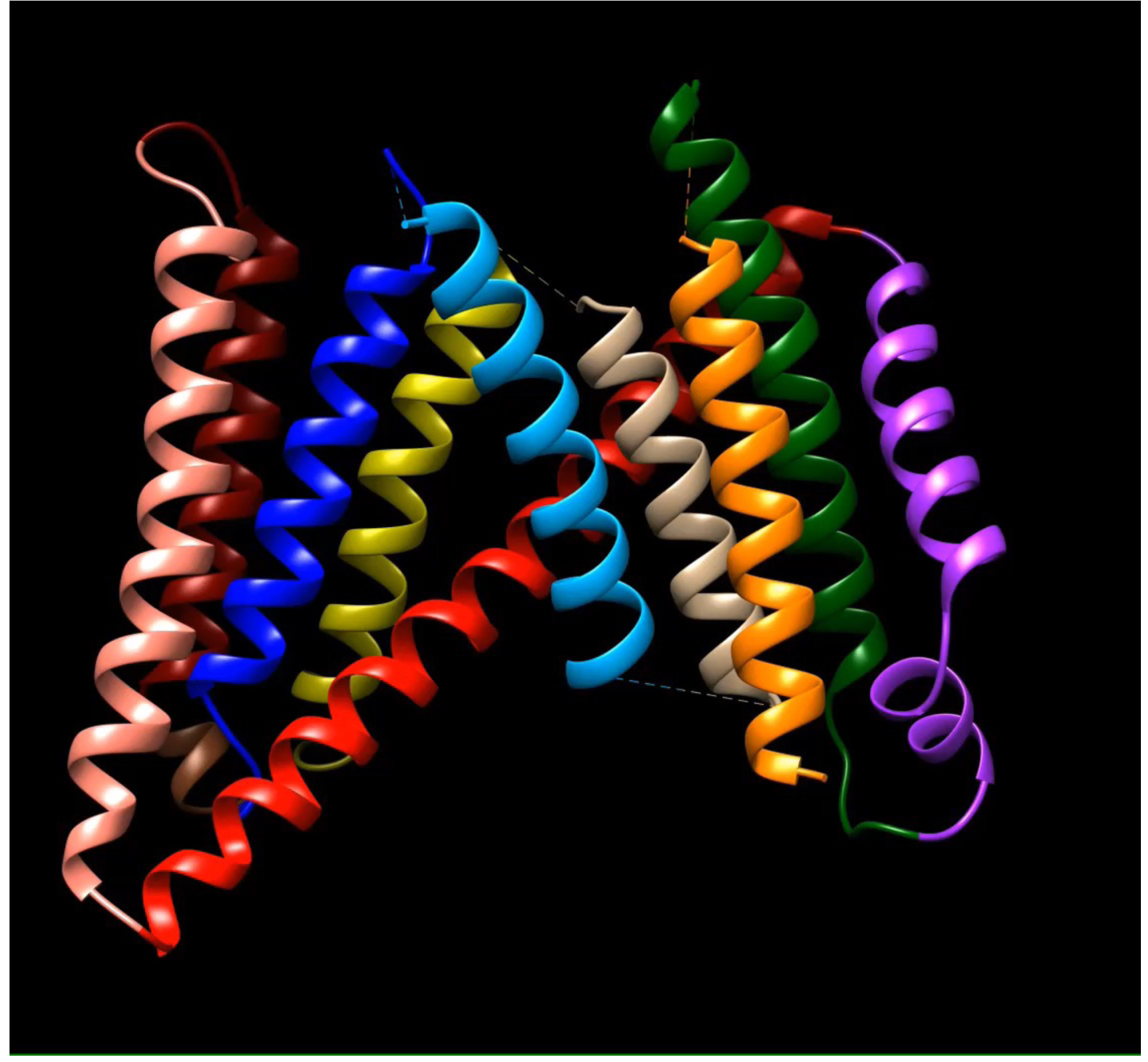
The top two SERINC5 AlphaFold models suggests a possible alternating access structural rearrangement. The movie shows the view from within the lipid bilayer of the conformations predicted by the top two models. The transmembrane *α*-helices are colored as in Figure 1e.

**Extended Data Movie 2.**
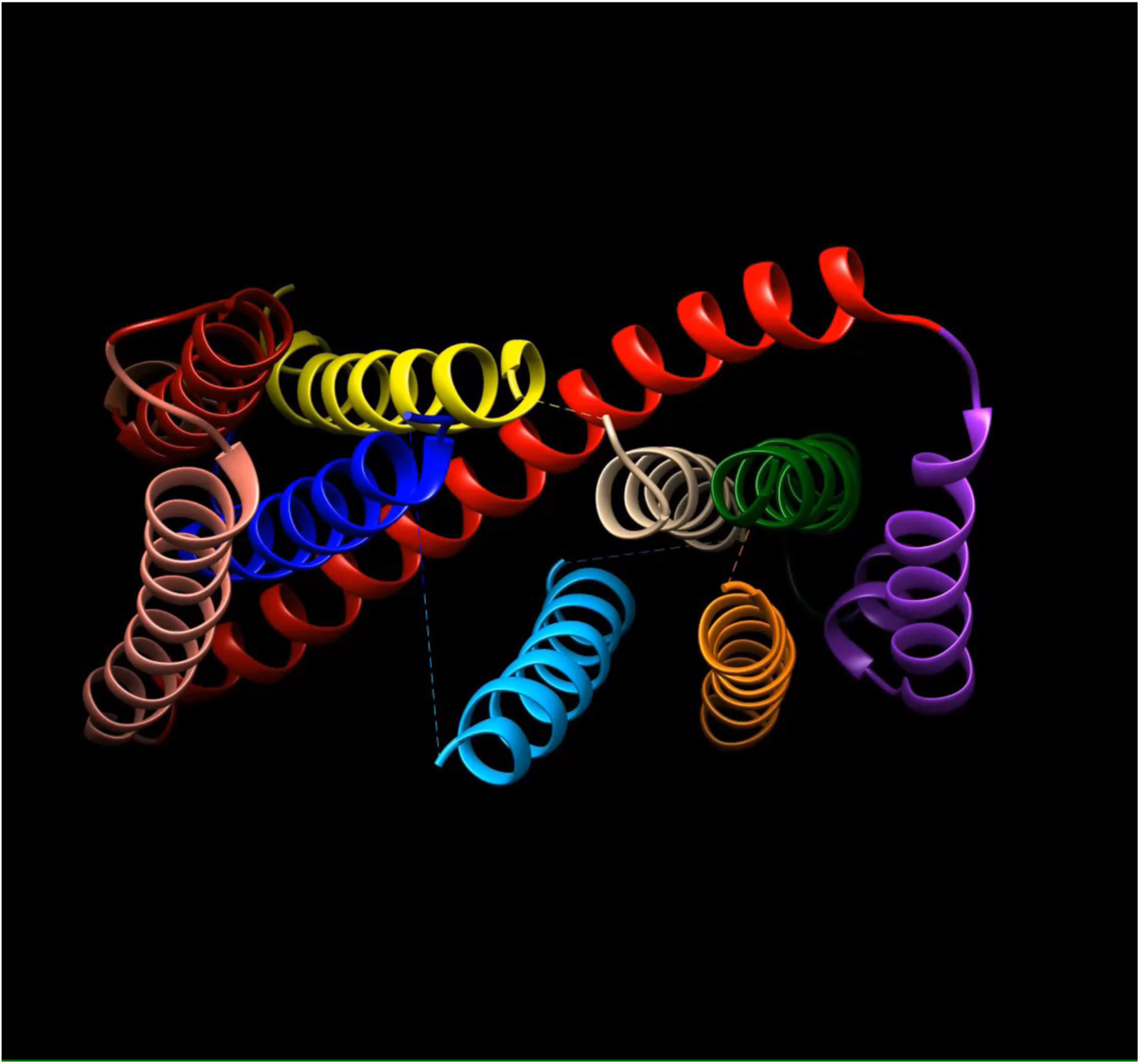
The top two SERINC5 AlphaFold models suggests a possible alternating access structural rearrangement. The movie shows the extracellular view of the conformations predicted by the top two models. The transmembrane *α*-helices are colored as in Figure 1e.

## References

1. Inuzuka, M., Hayakawa, M. & Ingi, T. Serinc, an activity-regulated protein family, incorporates serine into membrane lipid synthesis. J Biol Chem 280, 35776–35783, doi:10.1074/jbc.M505712200 (2005).

2. Rosa, A. et al. HIV-1 Nef promotes infection by excluding SERINC5 from virion incorporation. Nature 526, 212–217, doi:10.1038/nature15399 (2015).

3. Usami, Y., Wu, Y. & Gӧttlinger, H. G. SERINC3 and SERINC5 restrict HIV-1 infectivity and are counteracted by Nef. Nature 526, 218–223, doi:10.1038/nature15400 (2015).

4. Dai, W., Usami, Y., Wu, Y. & Gӧttlinger, H. A Long Cytoplasmic Loop Governs the Sensitivity of the Anti-viral Host Protein SERINC5 to HIV-1 Nef. Cell Rep 22, 869–875, doi:10.1016/j.celrep.2017.12.082 (2018).

5. Diehl, W. E. et al. Influence of Different Glycoproteins and of the Virion Core on SERINC5 Antiviral Activity. Viruses 13, doi:10.3390/v13071279 (2021).

6. Ahi, Y. S. et al. Functional Interplay Between Murine Leukemia Virus Glycogag, Serinc5, and Surface Glycoprotein Governs Virus Entry, with Opposite Effects on Gammaretroviral and Ebolavirus Glycoproteins. mBio 7, doi:10.1128/mBio.01985-16 (2016).

7. Ward, A. E. et al. HIV-cell membrane fusion intermediates are restricted by Serincs as revealed by cryo-electron and TIRF microscopy. J Biol Chem 295, 15183–15195, doi:10.1074/jbc.RA120.014466 (2020).

8. Chen, Y. C. et al. Super-Resolution Fluorescence Imaging Reveals That Serine Incorporator Protein 5 Inhibits Human Immunodeficiency Virus Fusion by Disrupting Envelope Glycoprotein Clusters. ACS Nano 14, 10929–10943, doi:10.1021/acsnano.0c02699 (2020).

9. Lai, R. P. et al. Nef decreases HIV-1 sensitivity to neutralizing antibodies that target the membrane-proximal external region of TMgp41. PLoS Pathog 7, e1002442, doi:10.1371/journal.ppat.1002442 (2011).

10. Sood, C., Marin, M., Chande, A., Pizzato, M. & Melikyan, G. B. SERINC5 protein inhibits HIV-1 fusion pore formation by promoting functional inactivation of envelope glycoproteins. J Biol Chem 292, 6014–6026, doi:10.1074/jbc.M117.777714 (2017).

11. Zhang, X. et al. CD4 Expression and Env Conformation Are Critical for HIV-1 Restriction by SERINC5. J Virol 93, e00544, doi:10.1128/JVI.00544-19 (2019).

12. Trautz, B. et al. The host-cell restriction factor SERINC5 restricts HIV-1 infectivity without altering the lipid composition and organization of viral particles. J Biol Chem 292, 13702–13713, doi:10.1074/jbc.M117.797332 (2017).

13. Stoneham, C. A. et al. A Conserved Acidic-Cluster Motif in SERINC5 Confers Partial Resistance to Antagonism by HIV-1 Nef. J Virol 94, e01554, doi:10.1128/JVI.01554-19 (2020).

14. Ren, X., Park, S. Y., Bonifacino, J. S. & Hurley, J. H. How HIV-1 Nef hijacks the AP-2 clathrin adaptor to downregulate CD4. Elife 3, e01754, doi:10.7554/eLife.01754 (2014).

15. Shi, J. et al. HIV-1 Nef Antagonizes SERINC5 Restriction by Downregulation of SERINC5 via the Endosome/Lysosome System. J Virol 92, e00196, doi:10.1128/JVI.00196-18 (2018).

16. Pye, V. E. et al. A bipartite structural organization defines the SERINC family of HIV-1 restriction factors. Nat Struct Mol Biol 27, 78–83, doi:10.1038/s41594-019-0357-0 (2020).

17. Sharom, F. J. Flipping and flopping--lipids on the move. IUBMB Life 63, 736–746, doi:10.1002/iub.515 (2011).

18. Sanyal, S. & Menon, A. K. Flipping lipids: why an’ what’s the reason for? ACS Chem Biol 4, 895–909, doi:10.1021/cb900163d (2009).

19. Contreras, F. X., Sanchez-Magraner, L., Alonso, A. & Goni, F. M. Transbilayer (flip-flop) lipid motion and lipid scrambling in membranes. FEBS Lett 584, 1779–1786, doi:10.1016/j.febslet.2009.12.049 (2010).

20. Pomorski, T. G. & Menon, A. K. Lipid somersaults: Uncovering the mechanisms of protein-mediated lipid flipping. Prog Lipid Res 64, 69–84, doi:10.1016/j.plipres.2016.08.003 (2016).

21. Zakrzewska, S. et al. Inward-facing conformation of a multidrug resistance MATE family transporter. Proc Natl Acad Sci U S A 116, 12275–12284, doi:10.1073/pnas.1904210116 (2019).

22. Kuk, A. C., Mashalidis, E. H. & Lee, S. Y. Crystal structure of the MOP flippase MurJ in an inward-facing conformation. Nat Struct Mol Biol 24, 171–176, doi:10.1038/nsmb.3346 (2017).

23. Kuk, A. C. Y., Hao, A., Guan, Z. & Lee, S. Y. Visualizing conformation transitions of the Lipid II flippase MurJ. Nat Commun 10, 1736, doi:10.1038/s41467-019-09658-0 (2019).

24. Zhang, B. et al. Structure of a proton-dependent lipid transporter involved in lipoteichoic acids biosynthesis. Nat Struct Mol Biol 27, 561–569, doi:10.1038/s41594-020-0425-5 (2020).

25. Nakano, M. et al. Flip-flop of phospholipids in vesicles: kinetic analysis with time-resolved small-angle neutron scattering. J Phys Chem B 113, 6745–6748, doi:10.1021/jp900913w (2009).

26. Homan, R. & Pownall, H. J. Transbilayer diffusion of phospholipids: dependence on headgroup structure and acyl chain length. Biochim Biophys Acta 938, 155–166, doi:10.1016/0005-2736(88)90155-1 (1988).

27. Ploier, B. & Menon, A. K. A Fluorescence-based Assay of Phospholipid Scramblase Activity. J Vis Exp, e54635, doi:10.3791/54635 (2016).

28. Goren, M. A. et al. Constitutive phospholipid scramblase activity of a G protein-coupled receptor. Nat Commun 5, 5115, doi:10.1038/ncomms6115 (2014).

29. Menon, I. et al. Opsin is a phospholipid flippase. Curr Biol 21, 149–153, doi:10.1016/j.cub.2010.12.031 (2011).

30. Bevers, E. M. & Williamson, P. L. Getting to the Outer Leaflet: Physiology of Phosphatidylserine Exposure at the Plasma Membrane. Physiol Rev 96, 605–645, doi:10.1152/physrev.00020.2015 (2016).

31. Schulte, B. et al. Localization to detergent-resistant membranes and HIV-1 core entry inhibition correlate with HIV-1 restriction by SERINC5. Virology 515, 52–65, doi:10.1016/j.virol.2017.12.005 (2018).

32. Suzuki, J., Umeda, M., Sims, P. J. & Nagata, S. Calcium-dependent phospholipid scrambling by TMEM16F. Nature 468, 834–838, doi:10.1038/nature09583 (2010).

33. Hankins, H. M., Baldridge, R. D., Xu, P. & Graham, T. R. Role of flippases, scramblases and transfer proteins in phosphatidylserine subcellular distribution. Traffic 16, 35–47, doi:10.1111/tra.12233 (2015).

34. Zaitseva, E. et al. Fusion Stage of HIV-1 Entry Depends on Virus-Induced Cell Surface Exposure of Phosphatidylserine. Cell Host Microbe 22, 99–110 e117, doi:10.1016/j.chom.2017.06.012 (2017).

35. Segawa, K., Suzuki, J. & Nagata, S. Constitutive exposure of phosphatidylserine on viable cells. Proc Natl Acad Sci U S A 108, 19246–19251, doi:10.1073/pnas.1114799108 (2011).

36. Le, T. et al. An inner activation gate controls TMEM16F phospholipid scrambling. Nat Commun 10, 1846, doi:10.1038/s41467-019-09778-7 (2019).

37. Munro, J. B. et al. Conformational dynamics of single HIV-1 envelope trimers on the surface of native virions. Science 346, 759–763, doi:10.1126/science.1254426 (2014).

38. Ma, X. et al. HIV-1 Env trimer opens through an asymmetric intermediate in which individual protomers adopt distinct conformations. Elife 7, doi:10.7554/eLife.34271 (2018).

39. Ding, S. et al. CD4 Incorporation into HIV-1 Viral Particles Exposes Envelope Epitopes Recognized by CD4-Induced Antibodies. J Virol 93, doi:10.1128/JVI.01403-19 (2019).

40. Yamaji-Hasegawa, A. & Tsujimoto, M. Asymmetric distribution of phospholipids in biomembranes. Biol Pharm Bull 29, 1547–1553, doi:10.1248/bpb.29.1547 (2006).

41. Lorent, J. H. et al. Plasma membranes are asymmetric in lipid unsaturation, packing and protein shape. Nat Chem Biol 16, 644–652, doi:10.1038/s41589-020-0529-6 (2020).

42. Salimi, H. et al. The lipid membrane of HIV-1 stabilizes the viral envelope glycoproteins and modulates their sensitivity to antibody neutralization. J Biol Chem 295, 348–362, doi:10.1074/jbc.RA119.009481 (2020).

43. Li, G. et al. The dual role of tetraspanin CD63 in HIV-1 replication. Virol J 11, 23, doi:10.1186/1743-422X-11-23 (2014).

44. Mucksch, F. et al. Quantification of phosphoinositides reveals strong enrichment of PIP2 in HIV-1 compared to producer cell membranes. Sci Rep 9, 17661, doi:10.1038/s41598-019-53939-z (2019).

45. Butan, C., Lara-Tejero, M., Li, W., Liu, J. & Galan, J. E. High-resolution view of the type III secretion export apparatus in situ reveals membrane remodeling and a secretion pathway. Proc Natl Acad Sci U S A 116, 24786–24795, doi:10.1073/pnas.1916331116 (2019).

46. Wu, X. et al. Structural basis of ER-associated protein degradation mediated by the Hrd1 ubiquitin ligase complex. Science 368, eaaz2449, doi:10.1126/science.aaz2449 (2020).

47. Feng, S. et al. Cryo-EM Studies of TMEM16F Calcium-Activated Ion Channel Suggest Features Important for Lipid Scrambling. Cell Rep 28, 567–579 e564, doi:10.1016/j.celrep.2019.06.023 (2019).

48. Bethel, N. P. & Grabe, M. Atomistic insight into lipid translocation by a TMEM16 scramblase. Proc Natl Acad Sci U S A 113, 14049–14054, doi:10.1073/pnas.1607574113 (2016).

49. Butler, E. K., Davis, R. M., Bari, V., Nicholson, P. A. & Ruiz, N. Structure-function analysis of MurJ reveals a solvent-exposed cavity containing residues essential for peptidoglycan biogenesis in Escherichia coli. J Bacteriol 195, 4639–4649, doi:10.1128/JB.00731-13 (2013).

50. Alvadia, C. et al. Cryo-EM structures and functional characterization of the murine lipid scramblase TMEM16F. Elife 8, e44365, doi:10.7554/eLife.44365 (2019).

51. Maxfield, F. R. & van Meer, G. Cholesterol, the central lipid of mammalian cells. Curr Opin Cell Biol 22, 422–429, doi:10.1016/j.ceb.2010.05.004 (2010).

52. Kumar, S., Rubino, F. A., Mendoza, A. G. & Ruiz, N. The bacterial lipid II flippase MurJ functions by an alternating-access mechanism. J Biol Chem 294, 981–990, doi:10.1074/jbc.RA118.006099 (2019).

53. Falzone, M. E. et al. TMEM16 scramblases thin the membrane to enable lipid scrambling. Nat Commun 13, 2604, doi:10.1038/s41467-022-30300-z (2022).

## References

1. Dominik, P. K. et al. Conformational Chaperones for Structural Studies of Membrane Proteins Using Antibody Phage Display with Nanodiscs. Structure 24, 300–309, doi:10.1016/j.str.2015.11.014 (2016).

2. Dominik, P. K. & Kossiakoff, A. A. Phage display selections for affinity reagents to membrane proteins in nanodiscs. Methods Enzymol 557, 219–245, doi:10.1016/bs.mie.2014.12.032 (2015).

3. Paduch, M. et al. Generating conformation-specific synthetic antibodies to trap proteins in selected functional states. Methods 60, 3–14, doi:10.1016/j.ymeth.2012.12.010 (2013).

4. Fellouse, F. A. et al. High-throughput generation of synthetic antibodies from highly functional minimalist phage-displayed libraries. J Mol Biol 373, 924–940, doi:10.1016/j.jmb.2007.08.005 (2007).

5. Fellouse, F. A., Wiesmann, C. & Sidhu, S. S. Synthetic antibodies from a four-amino- acid code: a dominant role for tyrosine in antigen recognition. Proc Natl Acad Sci U S A 101, 12467–12472, doi:10.1073/pnas.0401786101 (2004).

6. Zheng, S. Q. et al. MotionCor2: anisotropic correction of beam-induced motion for improved cryo-electron microscopy. Nat Methods 14, 331–332, doi:10.1038/nmeth.4193 (2017).

7. Zivanov, J. et al. New tools for automated high-resolution cryo-EM structure determination in RELION-3. Elife 7, e42166, doi:10.7554/eLife.42166 (2018).

8. Punjani, A., Rubinstein, J. L., Fleet, D. J. & Brubaker, M. A. cryoSPARC: algorithms for rapid unsupervised cryo-EM structure determination. Nat Methods 14, 290–296, doi:10.1038/nmeth.4169 (2017).

9. Grant, T., Rohou, A. & Grigorieff, N. cisTEM, user-friendly software for single-particle image processing. Elife 7, e35383, doi:10.7554/eLife.35383 (2018).

10. Emsley, P., Lohkamp, B., Scott, W. G. & Cowtan, K. Features and development of Coot. Acta Crystallogr D Biol Crystallogr 66, 486–501, doi:10.1107/S0907444910007493 (2010).

11. Pye, V. E. et al. A bipartite structural organization defines the SERINC family of HIV-1 restriction factors. Nat Struct Mol Biol 27, 78–83, doi:10.1038/s41594-019-0357-0 (2020).

12. Omasits, U., Ahrens, C. H., Muller, S. & Wollscheid, B. Protter: interactive protein feature visualization and integration with experimental proteomic data. Bioinformatics 30, 884–886, doi:10.1093/bioinformatics/btt607 (2014).

13. Jones, D. T. Protein secondary structure prediction based on position-specific scoring matrices. J Mol Biol 292, 195–202, doi:10.1006/jmbi.1999.3091 (1999).

14. Goddard, T. D. et al. UCSF ChimeraX: Meeting modern challenges in visualization and analysis. Protein Sci 27, 14–25, doi:10.1002/pro.3235 (2018).

15. Lomize, M. A., Pogozheva, I. D., Joo, H., Mosberg, H. I. & Lomize, A. L. OPM database and PPM web server: resources for positioning of proteins in membranes. Nucleic Acids Res 40, D370–376, doi:10.1093/nar/gkr703 (2012).

16. Kim, D. E., Chivian, D. & Baker, D. Protein structure prediction and analysis using the Robetta server. Nucleic Acids Res 32, W526–531, doi:10.1093/nar/gkh468 (2004).

17. Mukherjee, S. et al. Engineered synthetic antibodies as probes to quantify the energetic contributions of ligand binding to conformational changes in proteins. J Biol Chem 293, 2815–2828, doi:10.1074/jbc.RA117.000656 (2018).

18. Croll, T. I. ISOLDE: a physically realistic environment for model building into low- resolution electron-density maps. Acta Crystallogr D Struct Biol 74, 519–530, doi:10.1107/S2059798318002425 (2018).

19. Ploier, B. & Menon, A. K. A Fluorescence-based Assay of Phospholipid Scramblase Activity. J Vis Exp, e54635, doi:10.3791/54635 (2016).

20. Falzone, M. E. & Accardi, A. Reconstitution of Proteoliposomes for Phospholipid Scrambling and Nonselective Channel Assays. Methods Mol Biol 2127, 207–225, doi:10.1007/978-1-0716-0373-4_15 (2020).

21. Verchere, A. et al. Light-independent phospholipid scramblase activity of bacteriorhodopsin from Halobacterium salinarum. Sci Rep 7, 9522, doi:10.1038/s41598-017-09835-5 (2017).

22. McIntire, W. E., et al. CryoEM structure of the native A2A adenosine receptor-Gs alpha beta4gamma2 complex reveals two high affinity states and allosteric pathways linking GPCR Galpha and Gbeta: Implications for G protein subunit dissociation. submitted (2022).

23. Mulvania, T., Hayes, B. & Hedin, D. A flow cytometric assay for rapid, accurate determination of Baculovirus titers. BioProcess J 3, 47–53 (2004).

24. Jaakola, V. P. et al. The 2.6 angstrom crystal structure of a human A2A adenosine receptor bound to an antagonist. Science 322, 1211–1217, doi:10.1126/science.1164772 (2008).

25. Usami, Y., Wu, Y. & Gӧttlinger, H. G. SERINC3 and SERINC5 restrict HIV-1 infectivity and are counteracted by Nef. Nature 526, 218–223, doi:10.1038/nature15400 (2015).

26. Ahi, Y. S. et al. Functional Interplay Between Murine Leukemia Virus Glycogag, Serinc5, and Surface Glycoprotein Governs Virus Entry, with Opposite Effects on Gammaretroviral and Ebolavirus Glycoproteins. mBio 7, doi:10.1128/mBio.01985-16 (2016).

27. Munro, J. B. et al. Conformational dynamics of single HIV-1 envelope trimers on the surface of native virions. Science 346, 759–763, doi:10.1126/science.1254426 (2014).

28. Blott, E. J., Bossi, G., Clark, R., Zvelebil, M. & Griffiths, G. M. Fas ligand is targeted to secretory lysosomes via a proline-rich domain in its cytoplasmic tail. J Cell Sci 114, 2405–2416, doi:10.1242/jcs.114.13.2405 (2001).

29. Hankins, H. M., Baldridge, R. D., Xu, P. & Graham, T. R. Role of flippases, scramblases and transfer proteins in phosphatidylserine subcellular distribution. Traffic 16, 35–47, doi:10.1111/tra.12233 (2015).

30. Suzuki, J., Umeda, M., Sims, P. J. & Nagata, S. Calcium-dependent phospholipid scrambling by TMEM16F. Nature 468, 834–838, doi:10.1038/nature09583 (2010).

31. Segawa, K., Suzuki, J. & Nagata, S. Constitutive exposure of phosphatidylserine on viable cells. Proc Natl Acad Sci U S A 108, 19246–19251, doi:10.1073/pnas.1114799108 (2011).

32. Le, T. et al. An inner activation gate controls TMEM16F phospholipid scrambling. Nat Commun 10, 1846, doi:10.1038/s41467-019-09778-7 (2019).

33. Stirling, D. R. et al. CellProfiler 4: improvements in speed, utility and usability. BMC Bioinformatics 22, 433, doi:10.1186/s12859-021-04344-9 (2021).

34. Juette, M. F. et al. Single-molecule imaging of non-equilibrium molecular ensembles on the millisecond timescale. Nat Methods 13, 341–344, doi:10.1038/nmeth.3769 (2016).

35. Diehl, W. E. et al. Influence of Different Glycoproteins and of the Virion Core on SERINC5 Antiviral Activity. Viruses 13, doi:10.3390/v13071279 (2021).

36. Ding, S. et al. CD4 Incorporation into HIV-1 Viral Particles Exposes Envelope Epitopes Recognized by CD4-Induced Antibodies. J Virol 93, doi:10.1128/JVI.01403-19 (2019).

37. Brooks, B. R. et al. CHARMM: the biomolecular simulation program. J Comput Chem 30, 1545–1614, doi:10.1002/jcc.21287 (2009).

38. Mucksch, F. et al. Quantification of phosphoinositides reveals strong enrichment of PIP2 in HIV-1 compared to producer cell membranes. Sci Rep 9, 17661, doi:10.1038/s41598-019-53939-z (2019).

39. Scheres, S. H. & Chen, S. Prevention of overfitting in cryo-EM structure determination. Nat Methods 9, 853–854, doi:10.1038/nmeth.2115 (2012).

